# Graph-based tracing of dynamically functioning gene circuits in cell fate decisions with isoform resolution

**DOI:** 10.1101/2025.10.21.683820

**Authors:** Wanqi Li, Hao Dong, Zihan Zhou, Wenlin Fan, Yutong Zhou, Xiaochun Yang, Gang Xue, Zhiyuan Li

## Abstract

Transcription factor (TF) regulatory networks govern complex cellular processes, yet their inference from single-cell transcriptomes is hindered by TFs’ low abundance, tissue-specific functions, and isoform diversity. We evaluated the utility of transcription factor activity (TFA) to infer TF–TF interactions, demonstrating its superiority over expression-based methods in capturing tissue-specific, high-fidelity regulatory relationships. Using simulated and real-world transcriptomes across human hematopoiesis and early embryonic development, we validated TFA’s ability to render regulatory relationships. Through utilizing the concept of propagated regulome, we uncovered TFA’s utility in retrospectively resolving dynamic regulatory circuits. To address regulome limitations, we developed an embryonic stem cell-specific Consensus REgulome Database (CRED) with isoform resolution, revealing functional heterogeneity among TF isoforms. Applied to human preimplantation single-cell datasets, CRED outperformed existing regulomes, identifying isoform-specific regulators and tracing regulatory modules despite severe dropouts. Our work unleashes TFA as a robust tool for reconstructing TF networks and highlights the importance of transited, isoform-level, cell-type-specific analysis in unraveling transcriptional regulatory circuits.

**Highlights:**

1. TFA (Transcription Factor Activity) renders regulatory relationships with high fidelity compared to expression.
2. Using transited regulons enables TFA to retrospectively resolve dynamic regulatory interactions in cell fate decisions.
3. An integrated ESC-specific transited regulome database reveals isoform-level heterogeneity.
4. TFA successfully identifies key TF regulatory modules during human preimplantation from datasets with severe dropout.

## Introduction

Transcription factor (TF) regulatory networks orchestrate complex cellular behaviors, from developmental lineage specification to dynamic state transitions^1–3^. These networks, comprising direct and indirect interactions among TFs, form the core of gene regulation across biological systems^4–6^. For example, representative regulatory motifs such as cross-inhibition with self-activation (CIS) have been widely implicated in binary cell fate decisions^7^, while cascades of TF interactions guide embryonic development through sequential gene programs^8^. To understand these processes, systems biology has long sought to reconstruct TF regulatory networks from gene expression data^9–11^. Yet, the nature of TF regulation poses major challenges: (1) TFs function at the protein level, often as multimers^5,12,13^, with activities often regulated by post-translational modifications^14–16^. Consequently, assessing regulatory relationships solely based on TF mRNA abundance, i.e. expressions, is limited; (2) Technically, TF transcripts are often lowly expressed and noisy, suffering from severe dropout in single-cell data^17–19^. (3) Moreover, the snapshot nature of most transcriptomic assays fails to capture transient intermediates in cell fate transitions, obscuring the full regulatory landscape^20^. These limitations raise a central question: how can we move beyond expression profiles to accurately infer TF regulatory networks that govern cell dynamics?

To overcome the disconnection between TF expression and functional activity, the metric of transcription factor activity (TFA) has been developed to estimate TF functional states from the expression of their downstream targets, which are organized into regulons. TFA-based methods such as VIPER and DoRothEA have demonstrated broad utility across diverse biological contexts. They have been successfully used to nominate key regulators in cancer progression, drug response, and stem cell differentiation—situations where TF expression alone often fails to reflect regulatory influence^21,22^. Despite their success, most applications of TFA remain focused on identifying and comparing TF activities across conditions^23,24^. Meanwhile, their potential for inferring regulatory relationships among TFs themselves, i.e., reconstructing TF regulatory networks, has received little attention. Theoretically, TFA offers a promising avenue for this task: coordinated or causal shifts in TF activities may reflect underlying regulatory circuits or network motifs. However, no systematic framework or benchmarking effort has yet assessed the utility of TFA for mapping TF–TF interactions, leaving this dimension of its application largely unexplored.

Despite its promise, applying TFA to infer TF regulatory networks faces several critical challenges. First, there is a lack of systematic benchmarking to validate whether TFA truly improves upon expression-based approaches for reconstructing TF–TF interactions. While TFA offers a more functional readout of TF state, it remains unclear whether it better captures regulatory motifs embedded in complex systems. Furthermore, the influence of regulon composition—particularly the distinction between direct (e.g., ChIP-seq validated) and transitive (e.g., correlation-inferred) interactions—on TFA-based inference has not been rigorously evaluated.

Second, the cell-type specificity of regulome resources represents a major bottleneck. Currently established regulome databases primarily focus on collecting the most comprehensive libraries of regulatory interactions, often including those identified or inferred across various cell types and cancer cell lines^22,25,26^. However, TFs function in a highly tissue-dependent manner, indicating the gene sets regulated by TFs vary across cell types. This mismatch between the regulome source and the cell types may undermine the utility of the TFA metric, especially for inferring precise tissue-dependent TF–TF regulatory wiring.

Third, most TF-centric analyses operate at the gene symbol level, overlooking the functional complexity introduced by alternative splicing^27^. Recent studies revealed that isoforms of the same TF gene can differ in localization, binding specificity, and regulatory function^28^. For example, *POU5F1*, a TF responsible for embryonic stem cell maintenance, has isoforms with divergent capacities to sustain pluripotency^29,30^, while isoforms of another TF *MBD2* exhibit opposite roles in either silencing or activating *POU5F1*^27,31^. Collapsing multiple isoforms into a single gene-level entity may obscure their distinct regulatory roles and lead to misrepresentation of the underlying regulatory network. Therefore, isoform-level resolution is necessary to faithfully reconstruct TF regulatory interactions.

To this end, we systematically evaluated the feasibility of using TFA to infer TF–TF regulatory relationships. We began with both simulated networks and real-world spatial transcriptome with dropout events, demonstrating that TFA offers more consistent and high-fidelity regulatory relationships compared to expression profiles. We next applied the TFA to an 11-Node Hematopoiesis Network and recovered dynamically functioning gene circuits in endpoint cell populations, with results supported by a single-cell hematopoiesis atlas. Furthermore, we compared the use of direct and transited regulomes. Our analysis showed that transitive regulomes, despite being inferred, facilitated the identification of dynamically functioning TFs more effectively in complex developmental systems.

To investigate the influence of cell-type specificity in regulome and address the limitations of established regulome resources, we constructed an embryonic stem cell (ESC)-specific transited regulome database with isoform resolution. Based on that, we first quantified functional heterogeneity among TF isoforms from a regulon perspective, highlighting their orthogonality in regulatory functions. Finally, we showcased the utility of TFA in human preimplantation development, tracing isoform-level TF regulatory modules in single-cell datasets with severe dropouts. We can thereby propose new candidates for the fate-decision modules. Together, these efforts provide a comprehensive evaluation of TFA in reconstructing TF regulatory networks, from theoretical validation to practical implementation. Our integration of isoform-resolved regulomes paves the way for uncovering the fine architecture of transcript regulation beyond the gene level.

## Result

### Section 1: TFA outperforms expression in revealing TF regulatory relationships

TFA can be quantified by combining the expression changes of target genes to a TF and corresponding mode of regulation (activation/inhibition), i.e. regulon (**Figure** 1A)^21^. While TFA has been widely used to compare TF activity across different conditions, it remains unclear whether TFA could also be harnessed to characterize the regulatory relationships between TFs. In particular, we asked whether TFA-based correlations could more accurately reveal TF–TF interactions than TF expression-based correlations.

To explore this question, we constructed a bipotent gene regulatory network (hereinafter referred to as the Bipotent Network) designed to simulate a binary cell fate decision (**Figure** 1B), encompassing three fundamental stages of cell differentiation: 1) Cells exit the stem cell state, initiating lineage priming; 2) Cells specify one of two alternative fates, characterized by increased activity of either TF *X* or TF *Y*; 3) Cells commit to their chosen fate, activating downstream gene programs specific to each lineage (e.g., gene *X1*–*X5*). Based on the Bipotent Network, we then generated single-cell transcriptomic data using a transcriptomic data generator^32^ (see **Methods**). The resulting low-dimension embeddings clearly captured the progression from a stem-like state toward two mutually exclusive terminal fates (**Figure** 1C, **Figure** S1A). Within this system, a well-defined CIS regulatory motif governs the interaction between TFs *X* and *Y*, providing a benchmark for testing whether this relationship can be recovered through TFA.

We next compared the ability of gene expression and TFA to recover the cross-inhibition relationship between TFs *X* and *Y*. Based on gene expressions, their correlation appeared vague and inconsistent across pseudotime (**Figure** 1D, left). Of note, during early differentiation, *X* and *Y* even exhibited a transient positive correlation, contrary to the ground truth (**Figure** 1D, right). This misleading signal arose from the cascading propagation of the differentiation signals. Specifically, the upstream regulator *S3* simultaneously activated *X* and *Y*, masking their underlying antagonism. In contrast, the TFA method, even after we masked the mutual regulations between *X* and *Y* (**Figure** S1B, see **Methods**), captured a consistent and strong negative correlation between *X* and *Y* throughout the differentiation trajectory (**Figure** 1E, left), including in intermediate cell states where expression-based correlation failed (**Figure** 1E, right). To rule out artifacts from shared regulons between *X* and *Y,* we masked their overlapping regulatory targets in the commitment phase. The negative TFA correlation persisted (see **Methods**, **Figure** S1C), reinforcing that the interaction was driven by underlying network logic rather than database overlap. Overall, our observation suggests that TFA more faithfully reflects functional regulatory interactions, particularly in transitional states where gene expression is confounded by shared upstream signals.

To assess the applicability of TFA in identifying real-world regulatory relationships, we applied it to a spatial transcriptomic dataset from human embryonic heart development. This dataset, generated by Asp et al.^33^, profiles spatial transcriptomics across cardiac tissue sections at three developmental stages in the first trimester. Here, we focused on a representative sample at nine post-conception weeks, which includes annotated anatomical regions such as the ventricle, atrium, and outflow tract (**Figure** 1F). Within this dataset, we applied TFA with regulome from established databases^21,22^ and identified TF pairs with strongly anti-correlated activities across spatial locations (see **Methods**). One of the most representative examples is the *MESP1*–*MEOX2*: While their expression levels showed weak and noisy correlation (**Figure** 1G, left; **Figure** S1D), their inferred activities revealed a clear and robust negative relationship (**Figure** 1G, right), suggesting mutually exclusive regulatory roles. Spatially, high TFA of *MESP1* was observed in the ventricle region, while the outflow tract / large vessels showed high TFA of *MEXO2* (**Figure** 1H). Importantly, the regulons of *MESP1* and *MEOX2* share minimal target gene overlap (12/517 targets for *MESP1* and 12/1065 for *MEOX2*, **Figure** S1E), suggesting that their anti-correlated activities are not artifacts of regulon redundancy.

Although direct evidence that *MESP1* and *MEOX2* engage in mutual inhibition has not been experimentally explored, their TFA patterns and spatial distributions suggest functionally exclusive roles in early cardiac development. *MESP1* is a master regulator that up-regulates a group of markers of multiple cardiac cell types, including ventricular markers^34^. *MESP1* knock-in in human ESCs could facilitate cardiac ventricle development^35^, and *MESP1*-null embryos showed various anomalies of heart tube formation^36^. MEOX2, commonly known as a key regulator of transitions from myofibroblasts to fibroblasts^37^, is associated with senescence in vascular endothelial cells^38^ and highly expressed in cardiac endothelial cells^39^. Our analysis may point to a regulatory axis that is difficult to capture at the expression level but becomes evident through activity-based inference, inviting further investigation into the regulatory logic of early mesoderm patterning.

Together, these results demonstrate that TFA offers a more faithful readout of TF-regulatory interactions than expression profiles, both in computational simulations and in real biological systems. By capturing consistent activity patterns even in noisy or transitional cellular contexts, TFA enables the identification of functional antagonism between TFs that may be missed by conventional methods.

**Figure 1.**
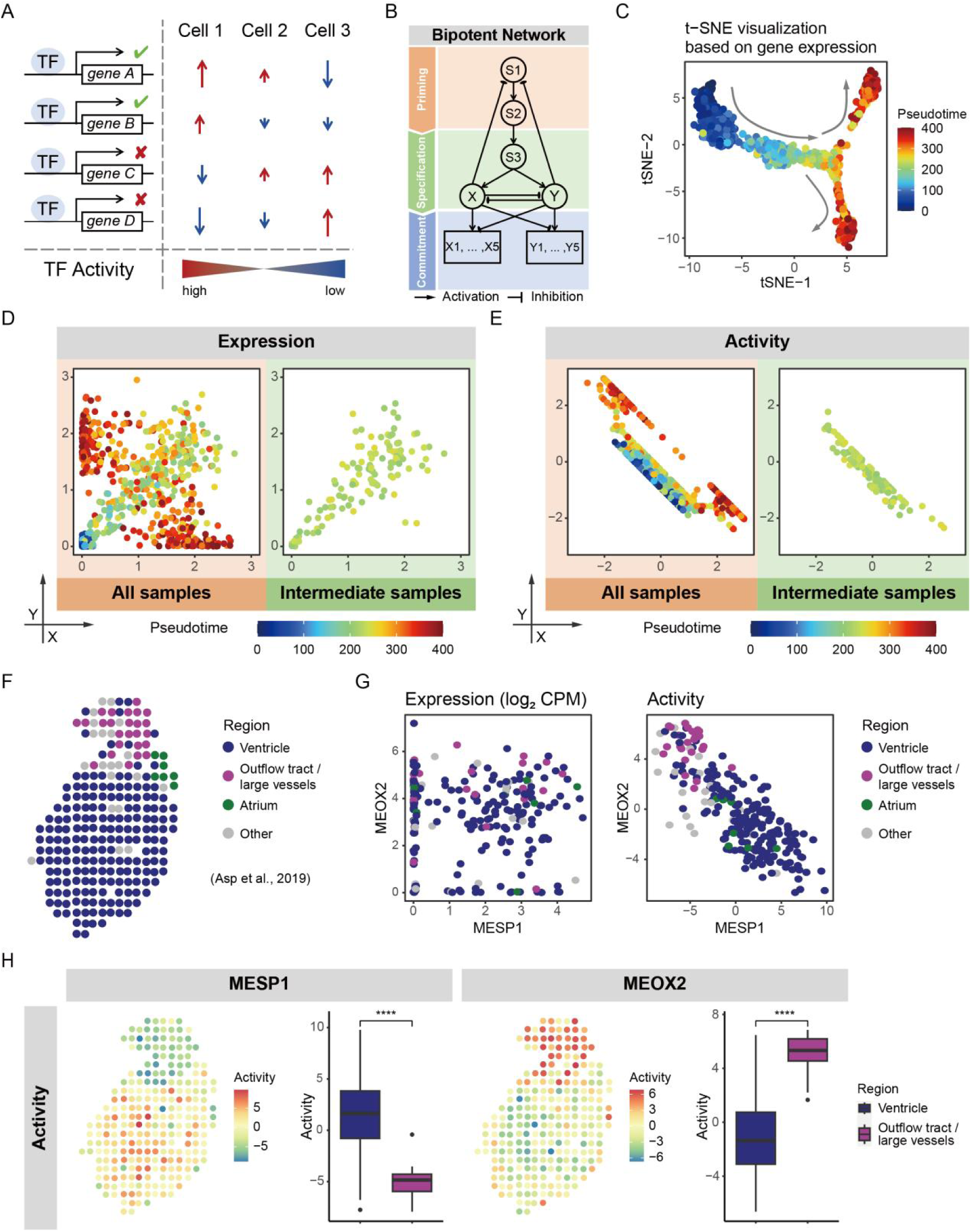
Comparison of TFA and gene expression in analyzing gene regulatory circuits. (A) Conceptual illustration of TFA. Green check marks indicate activating regulation, and red cross marks indicate repressing regulation. Upward red arrows and downward blue arrows represent increased and decreased gene expression, respectively. (B) Structure of the Bipotent Network with two antagonizing lineage-specifying factors *X* and *Y*. Source data of (C-E) is simulated result using this network. (C) T-SNE plot of simulated differentiation trajectories. Gray arrows indicate transiting directions over pseudotime. (D) Gene expression profiles of TFs *X* and *Y*. Intermediate samples are in the range of 200-250 pseudotime, corresponding to lineage specification stage. (E) TFA profiles of TFs *X* and *Y*. Intermediate samples are in the range of 200-250 pseudotime, corresponding to lineage specification stage. (F) Spatial transcriptomic data from a heart tissue section at 9 post-conception weeks (data from Asp et al.^33^). Color indicates anatomical region. (G) Gene expression (left) and TFA (right) of *MESP1* and *MEOX2* across spatial locations. (H) Activities of *MESP1* and *MEOX2* in spatial position and anatomical region. ∗∗∗∗ , p < 0.0001.

### Section 2: TFA enables retrospective inference of TF circuits from endpoint cell populations

A common limitation of single-cell transcriptomic datasets is their uneven sampling across differentiation trajectories. Intermediate or transitional cell states are often underrepresented, while terminally differentiated populations—referred to here as endpoint cell populations—tend to dominate^17^. This imbalance complicates efforts to reconstruct gene regulatory circuits that function dynamically during development. We therefore investigated whether TFA could be utilized to retrospectively trace the activity of upstream regulatory modules from the endpoint cells alone.

To explore this, we used an 11-node GRN for myeloid differentiation in hematopoiesis, established by Krumsiek et al. (**Figure** 2A; hereinafter referred to as the 11-Node Hematopoiesis Network)^40^. This network centers on mutual inhibition between *Gata1* and *Pu.1*, a canonical CIS fate-decision module in hematopoietic stem and progenitor cells (HSPCs)^41^. We simulated single-cell transcriptomic data based on this network, capturing trajectories where HSPCs eventually differentiated into four endpoint cell types (megakaryocyte, erythrocyte, monocyte, and granulocyte; **Figure** 2B-C, **Figure** S2A; see **Methods**). Focusing on granulocytes, the most abundant endpoint type in the simulation, we observed the expression pattern of highly expressed *Pu.1* and low expressed *Gata1* (**Figure** 2C-D). In this cell population, the expression profiles of the *Gata1*–*Pu.1* circuit exhibited negligible negative correlation (**Figure** 2D left), likely because the *Gata1*–*Pu.1* circuit primarily functions in upstream lineage bifurcation, not within granulocytes themselves (**Figure** S2B). In contrast, TFA recovered a strong negative relationship between *Gata1* and *Pu.1* within granulocytes (**Figure** 2E), suggesting that TFA patterns can preserve information about upstream regulatory dynamics, even after cells have committed to a terminal fate (**Figure** 2E).

We next validated this finding using a single-cell RNA-seq atlas of human hematopoiesis (**Figure** 2F and **Figure** S2C)^42^, with TFA calculated by regulome from established database^21,22^. Erythrocyte, the dominating endpoint cell type in this dataset, showed high *GATA1* expression but little to no detectable *SPI1* (also known as PU.1) expression due to dropout (**Figure** 2G, **Figure** S2D). Consistent with what we observed from the 11-Node Hematopoiesis Network simulated data, expression-based correlation between *GATA1* and *SPI1* was uninformative. In contrast, TFA successfully revealed a strong negative correlation between these two factors in erythrocytes (**Figure** 2H). This relationship could not be attributed to regulon overlap (**Figure** S2E), indicating that TFA captured a genuine signal of upstream antagonistic regulation. Notably, it has been recognized that the *GATA1*–*SPI1* circuit implements the fate choice between common myeloid progenitors (CMPs) and megakaryocyte-erythrocyte progenitor (MEPs), not in erythrocytes population^43^. Together, these results suggest that TFA retains the history of upstream regulatory events in endpoint populations, enabling retrospective characterization of dynamically functioning TF circuits even when intermediate states are under-sampled. This property makes TFA a valuable tool for decoding developmental logic from incomplete or imbalanced single-cell datasets.

**Figure 2.**
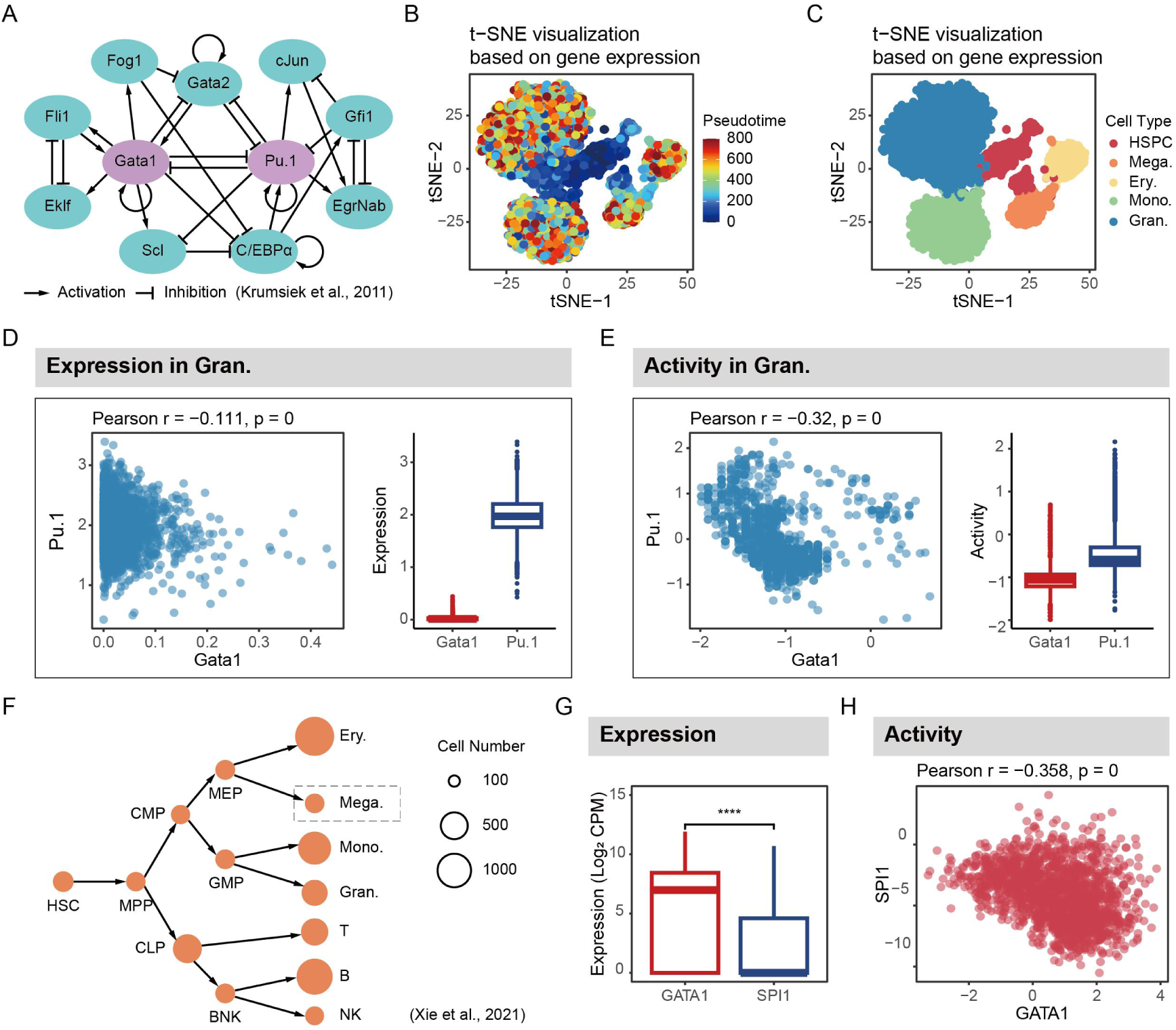
TFA reveals upstream regulatory circuits from endpoint cell populations. (A) A classical regulatory model of mouse myeloid differentiation from Krumsiek et al.^40^, highlighting the mutual inhibition between *Gata1* and *Pu.1* (purple) that were analyzed in (D) and (E). Source data of (B-E) is simulated result using this network. (B) T-SNE plot of simulated samples colored by pseudotime. (C) T-SNE plot of simulated samples colored by annotated cell types: HSPC, hematopoietic stem / progenitor cell; Mega., megakaryocyte; Ery., erythrocyte; Mono., monocyte; Gran., granulocyte. (D) Expression of *GATA1* and *PU.1* in granulocytes. Left: scatter plot of expressions; right: boxplot of expressions. (E) TFA of *GATA1* and *PU.1* in granulocytes. Left: scatter plot of activities; right: boxplot of activities. (F) Overview of lineages in human hematopoiesis, source data of (G) and (H). Circle size indicates cell number in Human blood cell atlas^42^. Cell type in gray frame line was absent in this dataset. HSC, hematopoietic stem cell; MPP, multipotent progenitor; CMP, common myeloid progenitors; MEP, megakaryocyte-erythrocyte progenitor; GMP, granulocyte-monocyte progenitor; CLP, common lymphoid progenitor. (G) Expression of *GATA1* and *SPI1* in erythrocytes. ∗∗∗∗, p < 0.0001. (H) TFA of *GATA1* and *SPI1* in erythrocytes.

### Section 3: Transited regulome facilitates the identification of master regulators

While the preceding results demonstrate that TFA has an advantage over gene expression in inferring regulatory relationships, even from endpoint cell populations, the accuracy of TFA depends tightly on the quality and structure of the regulome databases used. In practice, most public regulome resources are constructed by aggregating evidence from diverse biological contexts and comprise a mixture of direct and indirect (or transited) regulatory interactions (Here, we define direct regulons as gene targets directly bound and transcriptionally regulated by a TF, typically supported by ChIP-seq or reporter assays. In contrast, transited regulons refer to genes indirectly influenced by a TF through network propagation, such as via downstream intermediates in the gene regulatory networks)^22^. Although direct regulons offer more mechanistic specificity, transited regulons may better reflect system-level influence, especially when master regulators act far upstream of observable transcriptomic changes. Whether and how these two types of regulons affect the performance of TFA remains an open question, particularly in the context of dynamic cell state transitions.

To this end, we constructed a simplified unipotent network (**Figure** 3A; hereinafter referred to as the Unipotent Network) that models a linear cell differentiation trajectory from a unipotent progenitor into a mature cell state. The network contains three upstream regulators (*S1*–*S3*), followed by two modules of terminal genes (*X1*, *X2*). (**Figure** 3B, **Figure** S3A-D). Here, we curated two sets of regulomes: one set consists of direct regulations, while the other set contains indirect regulations (denoted as transited regulations as shown in **Figure** 3A). We compared the TFA profiles of TF *S3* using both regulome types. When using the direct regulome, the TFA could more precisely indicate the time window where *S3* exerted its function (**Figure** 3C, **Figure** S3B). Meanwhile, the transited regulome enabled the identification of upstream functioning TFs in the endpoint cell population (**Figure** 3D). Given the under-sampled intermediate states in single-cell RNA-seq, transited regulomes better uncover master regulatory genes driving key modules, beyond merely identifying hub genes with direct regulons (**Figure** 3E).

**Figure 3.**
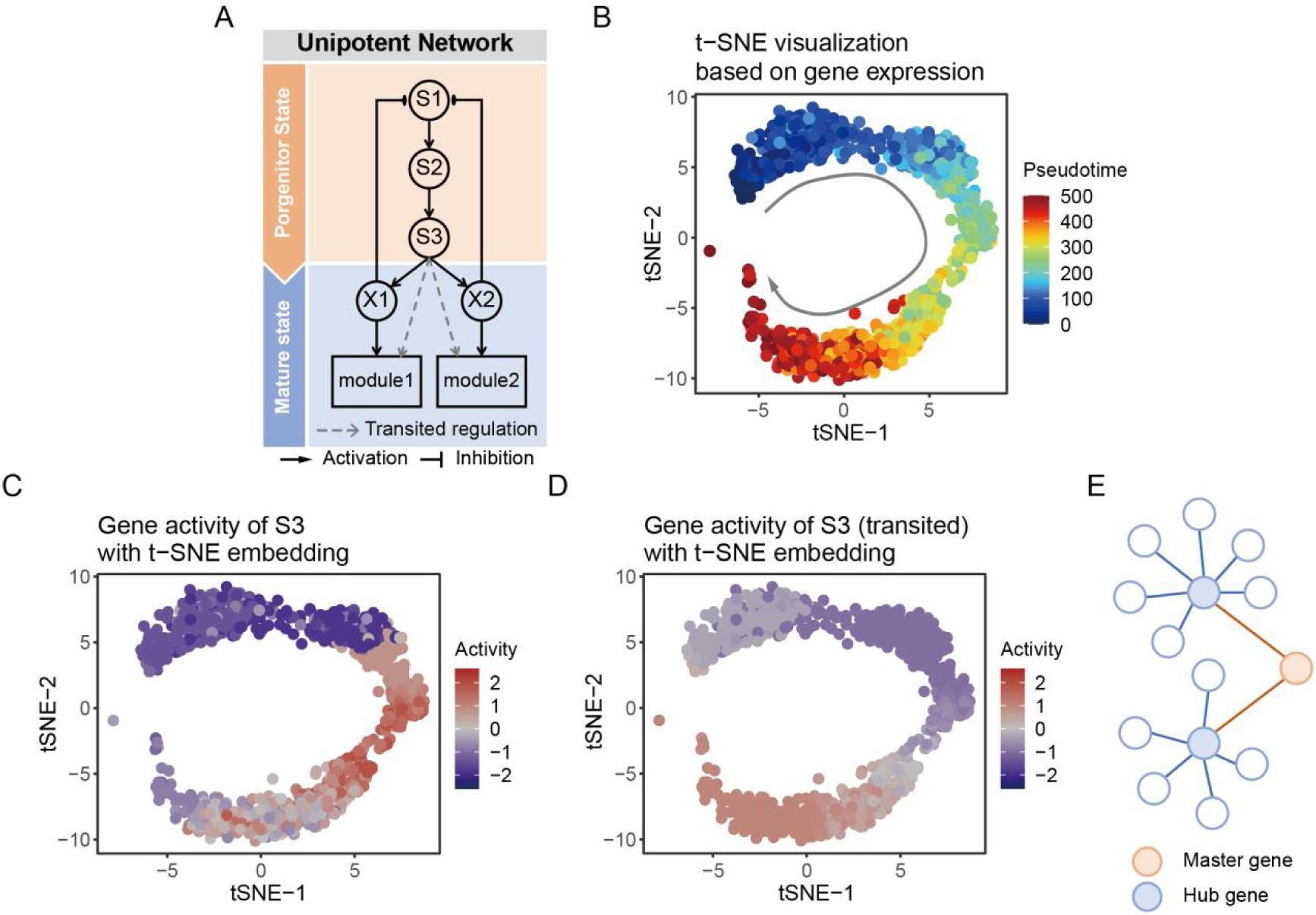
Transited regulome enables identification of master regulators from endpoint cells. (A) Structure of the Unipotent Network modeling the transition from progenitor to mature states: *S1*–*S3* form an upstream regulatory cascade; *S3* connects to two terminal modules (*X1* and *X2*) that inhibit the upstream module. Source data of (B-D) is simulated result using this network. (B) T-SNE plot of simulated trajectories. Gray arrows indicate transiting directions over pseudotime. (C) TFA of TF *S3* calculated using direct regulome. (D) TFA of TF *S3* calculated using transited regulome. (E) Conceptual diagram illustrating master regulators (orange) and hub genes (blue) in a regulatory network.

### Section 4: Integration of comprehensive regulome specific to embryonic development with isoform resolution

In previous sections, we demonstrated that TFA enables more accurate inference of TF–TF regulatory relationships compared to expression profiles, and transited regulome could help tracing the regulatory history from the endpoint cell population. However, the performance of TFA critically depends on the underlying regulome database. Given the complexity of multicellular organisms, two major limitations hinder established databases: (1) TFs function in a highly tissue-dependent manner, but most regulome databases are compiled across heterogeneous contexts. (2) Alternative splicing occur more frequently in TFs^44^, producing isoforms that can differ markedly in their regulatory functions, yet current interaction databases capture only a small fraction of known TF isoforms^28^. Thus, it remains unclear to what extent tissue-specific and isoform-level regulome impacts the inference of TF regulatory networks using TFA, and whether it is feasible to construct such a resource tailored for developmental contexts like ESCs.

Given their pluripotency, ESCs are widely used in broad researching fields like organoids and induced pluripotent stem cells (iPSCs), serving as starting point for development and reprogramming. Understanding ESC-specific regulatory programs is fundamental to advancing these fields. Here, we integrated a comprehensive human ESC-specific regulome database termed **C**urated **RE**gulome **D**atabase (CRED), with isoform-level resolution. We utilized a TF Atlas of directed differentiation that overexpressed 3,548 isoforms of 1,836 TFs respectively in human ESCs (hESCs)^1^, which enabled assessment of regulatory interactions with isoform resolution. In each TF isoform’s overexpression profile, we extracted up-regulated and down-regulated differentially expressed genes (DEGs) as its transited activation and inhibition targets, respectively (**Figure** 4A, see **Methods**). To reflect current understanding of regulatory network, where a notable proportion of TFs function as high-connectivity hubs, we chose the cutoff of DEGs that yield out-degree distribution most consistent with existing databases (TRRUST^25,26^, DoRothEA^22^, **Figure** S4A, B). The resulting CRED contains 1,180,085 interactions of 3,250 TF isoforms, while most interactions were activation (1,130,532, 95.80%). 24,350 proteins were involved in the GRN assembled by TF–gene interactions in CRED (**Table** S1). In benchmarking against widely used regulome databases (TRRUST^25,26^, DoRothEA^22^), we collapsed isoform interactions back to gene symbols (see **Methods**). We next investigated whether isoforms of the same TF share similar regulons in CRED. Our analysis revealed that only 31.09% of the interaction is shared, indicating that most isoforms exhibit distinct regulatory functions. This finding aligns with previous studies showing that 73% of gene–gene interactions differ across isoforms, supporting the concept that alternative splicing generates functionally diverse protein variants with specialized regulatory roles^28^.

The low overlap between databases highlights substantial tissue specificity of regulatory interactions (**Figure** S4D). To assess whether this specificity translates into improved performance, we benchmarked CRED in a dataset of combinatorial TF overexpression in hESCs from the project of TF Atlas^1^, where overexpressed TFs would activate their downstream targets and consequently rank highly in TFA. Comparative analysis of TFA calculation using different regulome databases (CRED, established database represented by TRRUST^25,26^ and DoRothEA^22^, and merged database) showed that CRED achieved the highest recall rate, followed by merged database (**Figure** 4B). These results confirm that context-specific regulome—rather than broad cross-tissue aggregates—are more effective for accurate TFA estimation, particularly in stem cell systems

Previous works have shown that TF isoforms can exhibit remarkable functional divergence, often exceeding that observed in paralogous TFs^28^. Functional divergence underscores the necessity of analyzing regulatory interactions at the isoform level. Using the comprehensive regulome covered in CRED, we quantified pair-wise regulatory similarity between isoforms by using Jaccard index, where higher value indicates greater proportion of shared regulons. The first observation is that most isoforms exhibited low regulatory similarity (Jaccard index < 0.3), necessitating isoform-level regulome. Next, hierarchical clustering identified a subgroup with substantially higher Jaccard indices, suggesting high functional coherence (**Figure** 4C, cluster 4). Cluster 4 contained 417 isoforms representing 283 TFs, significantly enriched for TFs with multiple isoforms (**Figure** 4D-E). Despite this functional coherence, variability still existed: some TFs had all isoforms grouped within Cluster 4, while others had isoforms both within and outside Cluster 4 (**Figure** S4E). However, a subset of 30 TFs, which we termed “identical TFs”, showed complete inclusion of all isoforms in cluster 4, indicating they shared highly conserved regulatory programs (**Figure** S4F). Gene ontology (GO) enrichment analysis revealed that these “identical TFs” predominantly govern core cellular programs, including transcriptional initiation, metabolic regulation, and homeostatic responses. While a subset also participates in early developmental and differentiation processes, the overall enrichment points to broadly required regulatory roles that may favor isoform conservation (**Figure** 4F).

In conclusion, we constructed an ESC-specific regulome CRED that outperformed existing databases in identifying active TFs within the ESC context. Notably, aggregating cross-tissue regulons impairs the TFA performance, underscoring the importance of tissue specificity. Moreover, the pronounced functional divergence among isoforms highlights the necessity of isoform-level resolution in accurately inferring gene regulatory relationships.

**Figure 4.**
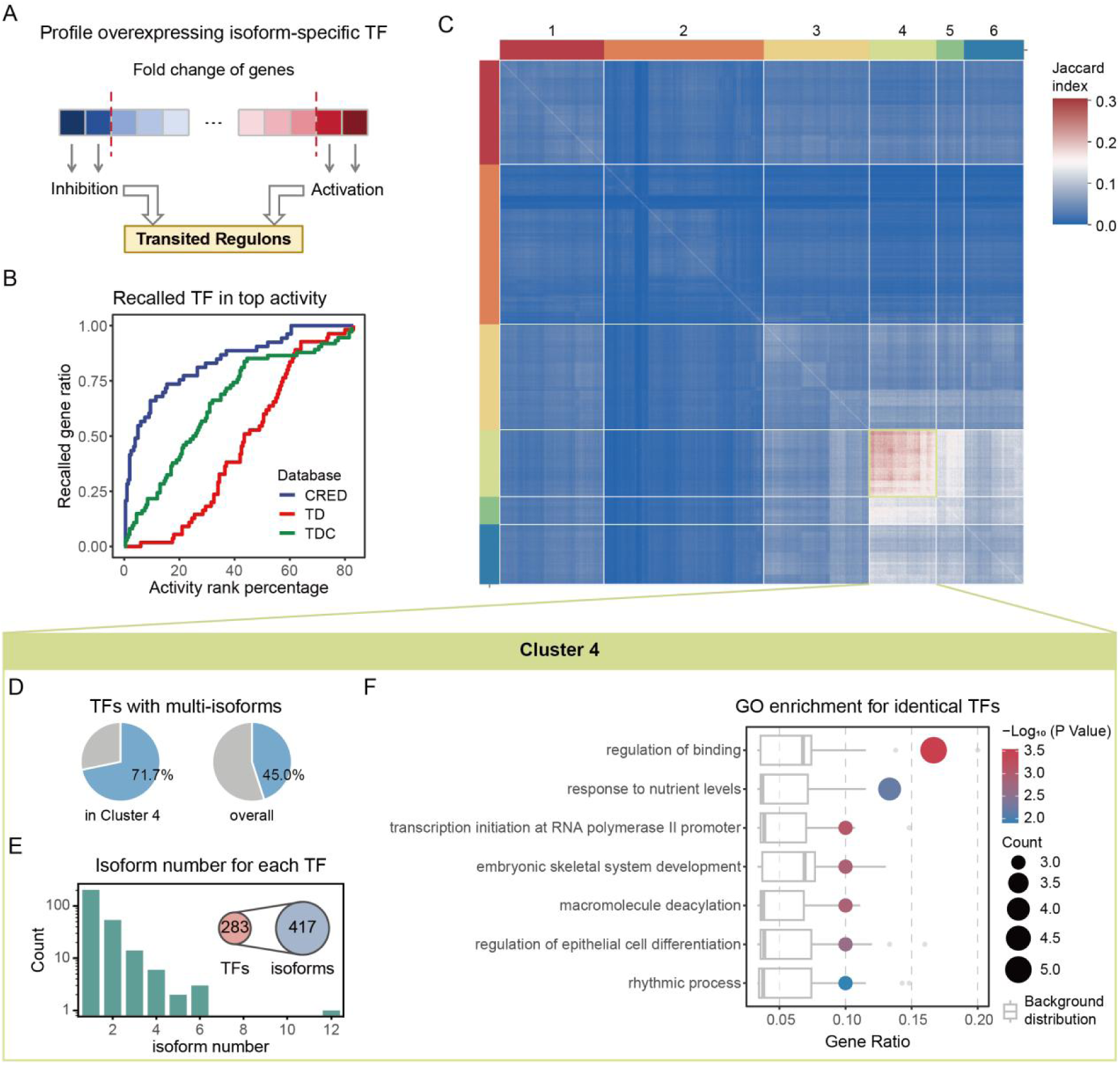
Construction, benchmarking, and clustering analysis of CRED. (A) Schematic workflow for generating transited ESC-specific regulome CRED from isoform-specific overexpression profiles. (B) Recall curve of detecting overexpressed TF combinations as high TFA using different regulome databases. TD, merged database using TRRUST^25,26^ and DoRothEA^22^. TDC, merged database using TRRUST^25,26^, DoRothEA^22^ and TF resolution CRED. (C) Hierarchical clustering of isoform regulons based on Jaccard index of regulations. (D-F) focused on Cluster 4 (light green square). (D) Ratio of TFs with multiple isoforms in Cluster 4 versus the full database. (E) Distribution of isoform number per TF in Cluster 4, with the total numbers of isoform and TFs involved in Cluster 4. (F) GO enrichment analysis for “identical TFs” in Cluster 4. Top terms of biological process with gene ratio higher than upper quartile of background distribution (generated by 50 random TF lists with same size) were listed.

### Section 5: Regulon-based analysis underlines orthogonality in function between isoforms

Given the substantial diversity in regulatory targets among TF isoforms in CRED, we next systematically characterized isoform-level regulatory divergence across all TFs and assessed their functional orthogonality. To quantify each isoform’s regulatory uniqueness, we defined “unique interactions” as those present in only one isoform (absent in all others) or with opposite signs in all other isoforms—essentially, interactions not shared identically by any other isoform of the same TF (**Figure** 5A). In CRED, most isoforms possessed 100-500 unique interactions, reflecting substantial specificity (**Figure** S5A).

To quantify the degree of functional specificity of each isoform, we introduced a metric called unique interaction ratio (UIR), defined as the proportion of unique interactions relative to total regulon number (**Figure** S5B; see **Methods**). To evaluate isoform heterogeneity of each TF, we calculated the mean and range of UIR from all its isoforms (**Figure** 5B). Mean UIR measures the average level of isoform orthogonality, and range measures the heterogeneity of orthogonality (**Figure** 5C). TFs with low mean and low range of UIR had largely functional redundant isoforms—resembling the “identical TFs” cluster defined in Section 4. Using this “identical TFs” cluster as a baseline, we qualitatively categorized TFs into four distinct patterns of isoform orthogonality (**Figure** 5C, **Figure** S5C). Most TFs gathered around the lower right corner, meaning all of their isoforms function in highly orthogonal ways, including *GATA4*, *MBD2* and *SALL4* which previous research had revealed their diversity in isoform function^27,31,45,46^. Some TFs with high range of UIR exhibited more complex pattern, particularly containing both redundant isoforms and outliers with exceptionally high UIR (upper left corner of **Figure** 5C**, Figure** S5D). Overall, the mean and range of UIR showed a stronger negative correlation compared with background distribution generated by random shuffle (**Figure** 5B, **Figure** S5E). This observation suggests that TFs with highly orthogonal isoforms tend to be consistently orthogonal, while TFs with redundant isoforms still preserve a few highly orthogonal isoforms – orthogonality in isoforms seems to be a general trend. Analysis of TFs with same number of isoforms excluded the effect of isoform number (**Figure** S5F).

Next, we tested whether isoform regulatory heterogeneity is associated with the function of the TF. Interestingly, TFs regulating house-keeping functions like proliferation or apoptosis did not show remarkably different UIR patterns compared to others (**Figure** 5D). Similarly, TFs related to early development and cell fate decisions also showed no significant shift in UIR distribution (**Figure** 5E), leading to the conclusion that isoform orthogonality is decoupled with their general functions or associated biological processes.

In summary, our graph-based analysis revealed that isoforms of the same TF often regulate distinct target sets, and such orthogonality is widespread across functional categories. This underscores the necessity of adopting isoform-level regulome frameworks, as gene-level annotations would overlook substantial regulatory diversity critical for accurate interpretation of transcriptional programs.

**Figure 5.**
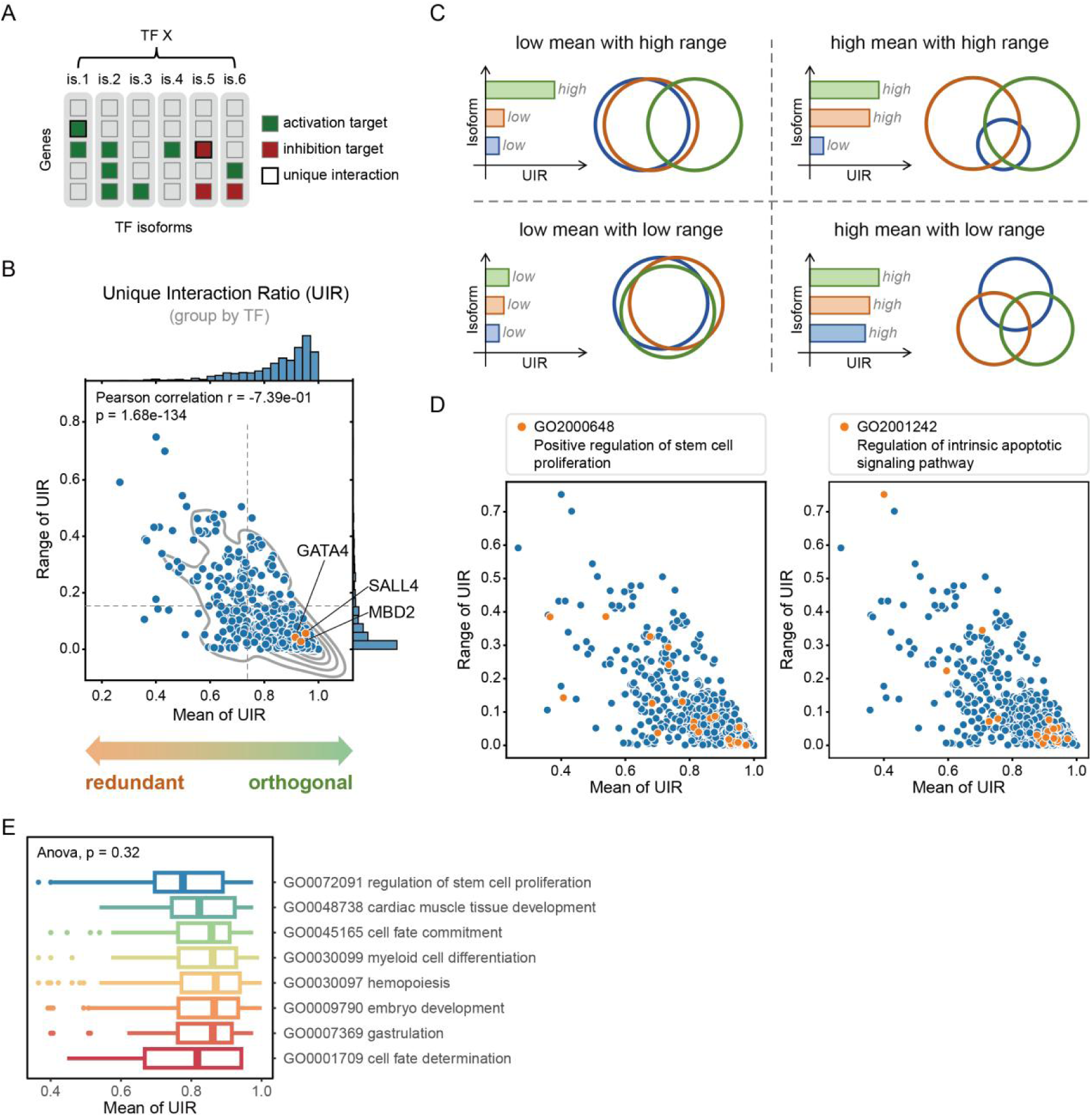
Regulon-based analysis reveals functional orthogonality among TF isoforms. (A) Schematic definition of isoform-unique interactions. For each isoform of a TF (is.n, no.n isoform of the TF), interactions not shared with other isoforms (by target gene or mode of regulation) are labeled as unique (highlighted in bold frame). (B) Mean and range of UIR grouped for each TF with more than one isoform. Each dot represents one TF. Orange dots highlight examples mentioned in the text. Dashed lines represent thresholds defined by “identical TFs” (see **Figure** S5C). (C) Conceptual diagram illustrating four typical scenarios of isoform orthogonality patterns based on mean and range of UIR. Each circle in the Venn diagrams represents the set of regulatory interactions for a given isoform. (D) Mean and range of UIR grouped by TFs involved in selected GO terms related to core biological functions: “positive regulation of stem cell proliferation” and “intrinsic apoptotic signaling pathway”, focusing on TFs in GO terms. (E) Distribution of mean UIR grouped by TF, focusing on TFs in GO terms related with development.

### Section 6: Application of isoform-level TFA reveals germ layer–specific regulators in human early development

To evaluate the practical utility of our TFA-based framework and the CRED regulome, we applied it to a real-world biological system mimicking post-implantation stages of human early embryo development, using a dataset of stem-cell-derived peri-gastruloids^47^. This in-vitro system captures key features of human gastrulation, where stem cells were prompted to self-organize and resulted in an embryo-like structure. In particular, single-cell RNA-seq data of day 11 peri-gastruloids could distinguish typical cell types in gastrula and here we focused on the three germ layers (**Figure** 6A, see **Methods**).

We first assessed whether CRED, our ESC-derived regulome database, could more accurately capture germ layer-associated TFA compared to conventional regulome databases. Given the limited availability of isoform-level annotations in other resources, we used the TF-level version of CRED for fair comparison. TFA calculated by CRED better reflected germ layer specificity: TFs known to be associated with a given layer showed significantly higher TFA in that lineage. Merged databases such as TRRUST^25,26^ and DoRothEA^22^ yielded noisier patterns (**Figure** 6B). Furthermore, we tested a curated set of 65 differentiation-inducing TFs identified in context of human iPSCs^48^, which should be highly active in the offspring of human iPSCs, e.g. gastruloids. CRED outperformed other regulome databases in recalling differentiation-inducing TFs as top ranks in TFA, while other databases failed to outperform random permutation (**Figure** 6C).

As shown in earlier sections, TFA enables the recovery of upstream fate-regulators from endpoint cells. We then testified a group of TFs reported to drive hESCs-to-endoderm differentiation^49,50^. CRED-based TFA of these TFs was significantly higher in endoderm cells, while TFA calculated with other databases was less consistent (**Figure** 6D). Among these important endoderm-driving TFs, *KLF8* is known to play a pivotal role in accelerating the differentiation of definitive endoderm cells^50^. Multiple isoforms of *KLF8* have been detected, but their individual roles in hESC-to-endoderm transitions remains unclear^51^. Due to severe dropout in *KLF8* expression (**Figure** 6E), it served as an ideal example of TFA-based investigations.

Using CRED, four isoforms of *KLF8* exhibited distinct TFA patterns (**Figure** 6F, G, **Figure** S6A). Among them, *KLF8-2* was the only isoform that had relatively high TFA in endoderm, suggesting it may be the dominating isoform driving endoderm differentiation. For cross-validation, we directly examined isoform expressions at earlier stage of development stage using an isoform-resolved dataset of human preimplantation embryo transcriptome^52^. Only two of the *KLF8* isoforms (*KLF8-2*, *KLF8-3*) were detectable, indicating that the other two were expressed below the sequencing threshold. Through embryonic developmental stages, *KLF8-2* showed a gradually but strongly increased expression especially in morula and blastocyst, while *KLF8-3* only slightly increased (**Figure** 6H). This dominant expression trajectory of *KLF8-2* suggests its functional relevance and aligns with our TFA-based inference, demonstrating the power of CRED to resolve isoform-specific TFA even from gene-level expression data.

Next, we focused on *POU5F1* and *NANOG*, two TFs crucial for maintaining stem cell pluripotency during embryonic development^8,53^. *POU5F1* exhibited low level of dropout (**Figure** 6I), making expression a reliable proxy for TFA. Among four *POU5F1* isoforms, *POU5F1-1* showed the highest correlation between TFA and expression (**Figure** 6J, K, **Figure** S6B). It was also the only isoform detected in the dataset of human preimplantation embryo and displayed a steady increase along development stages (**Figure** 6L), suggesting *POU5F1-1* is the dominant isoform during gastrulation. In contrast, *NANOG* showed severe dropout except in ectoderm (**Figure** 6M). Despite the similar pattern across three germ layers of TFA for the two isoforms (**Figure** S6C), *NANOG-1* and *NANOG-2* showed inconsistent TFA (**Figure** 6N) where *NANOG-2* aligned closely with TF expression (**Figure** S6D). However, in the isoform-resolved dataset of human preimplantation embryo, *NANOG-1* had consistently higher expression (**Figure** 6O), beginning as early as the 8-cell stage. Considering expression of *NANOG-2* showed slight but significant increase in blastocyst stage and expression of *NANOG-1* was quite stable from 8-cell to blastocyst stage, we might here suggest that an isoform-switching event occurred. Such switching event of other TFs such as *FOXP1*^54^, *REST*^55^, and *GRHL1*^56^ has been reported in key stages of development, and vast majority of TFs would switch in isoform proportion, more frequently in developmental context^28^. In this context, we suggest *NANOG-1* functions at earlier stage and *NANOG-2* switches to dominant since blastocyst.

**Figure 6.**
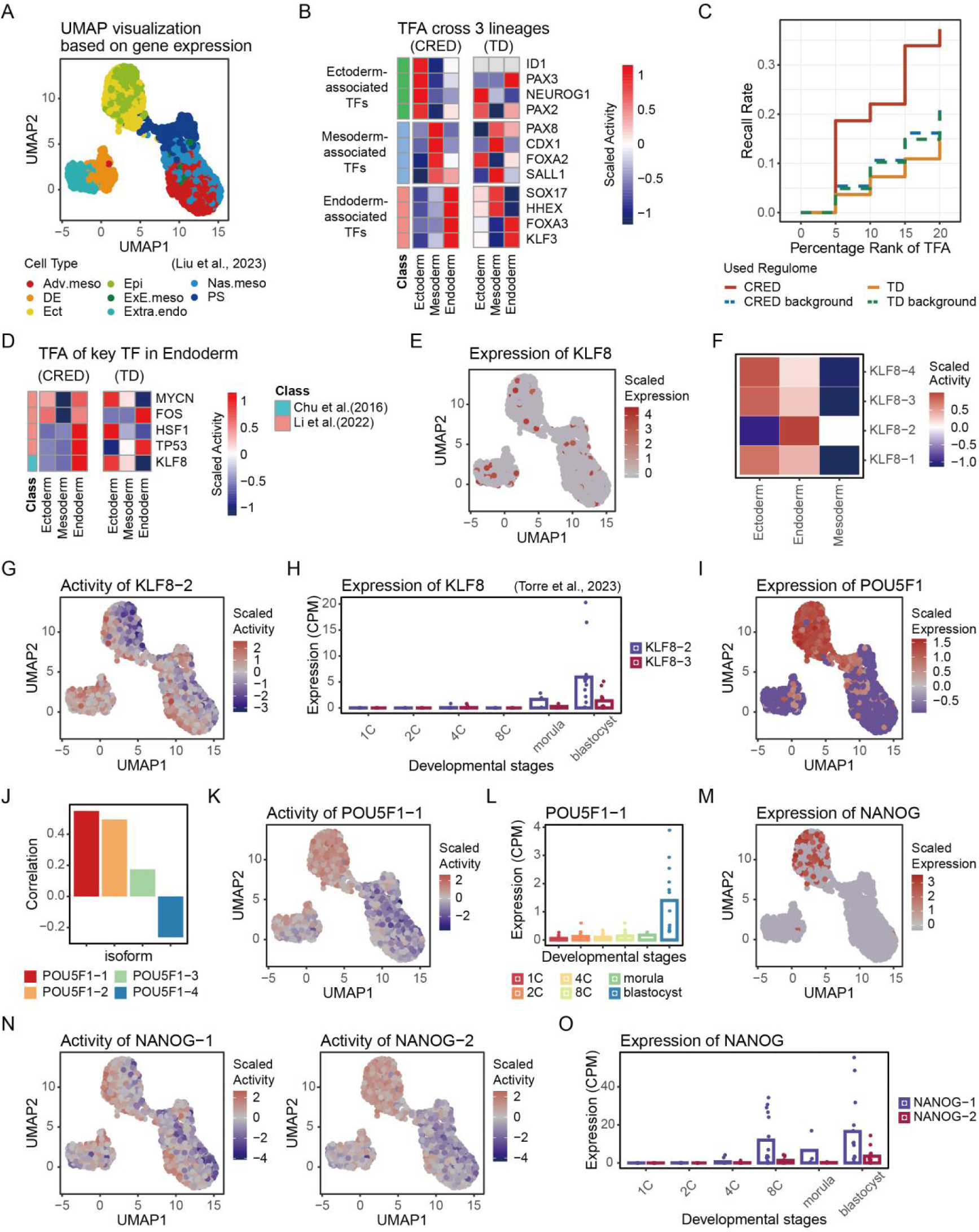
Isoform-level TFA in human early embryogenesis. (A) UMAP plot of human peri-gastruloids (Liu et al., 2023^47^, cells were downsampled), source data of (B-G), (I-K), (M) and (N) is this dataset. (B) TFA cross three lineages using different regulome database, focusing on lineage-associated TFs. (C) Recall rate of differentiation-inducing TFs^48^ in TFs with top TFA using different regulome databases. Background baselines were generated by random permutation. (D) TFA cross three lineages using different regulome databases, focusing on key TFs for endoderm development^49,50^. (E) Scaled expression of *KLF8*. (F) Scaled TFA of *KLF8* isoforms cross three lineages. (G) Scaled TFA of *KLF8-2*. (H) Expression of *KLF8* isoforms in human preimplantation embryo^52^, source data of (L) and (O) is also this dataset. 1C, zygote. 2C, 2-cell embryo. 4C, 4-cell embryo. 8C, 8-cell embryo. (I) Scaled expression of *POU5F1*. (J) Correlation of expression and TFA of *POU5F1* isoforms. (K) Scaled TFA of *POU5F1-1*. (L) Expression of *POU5F1-1* in human preimplantation embryo. (M) Scaled expression of *NANOG*. (N) Scaled TFA of *NANOG* isoforms. (O) Expression of *NANOG* isoforms in human preimplantation embryo.

### Section 7. Reconstruction of isoform-level gene circuits in early human development

To further explore the potential of our ESC-specific regulome database, we aimed to extract isoform-level gene circuits in human gastrulation by CRED. As we stressed in Section 2, TFA captured key regulators that dynamically functioned earlier, hence gene circuits we extracted with three germ layers of human peri-gastruloids should be responsible for the differentiation of ectoderm and mesoendoderm. This process could be modeled as bifurcation between ectoderm and mesoendoderm as antagonistic cell fates, therefore, TFs orchestrating these two fates would show a negative correlation in TFA, as we indicated in Section 1.

We therefore computed isoform-level TFA across the dataset of human peri-gastruloids, and then extracted top 2500 most negatively correlated isoform-level TF pairs in ectoderm cells and mesoendoderm cells, respectively. The 83 overlapping pairs across both lists were retained as putative cross-inhibition circuits in the whole process driving the bifurcation (**Figure** 7A). These 83 isoform pairs involved 58 TFs, among which 38 (65.52%) have been reported to regulate germ layer formation in human, mouse or other model organisms (**Figure** 7B). The remaining 20 TFs represent candidates of novel regulators for ectoderm-mesoendoderm fate decisions.

We next assembled all isoforms involved in the 83 pair into a network, with edges representing significant negative correlation. Of note, several isoforms of the same TF frequently appeared in network and often connected to similar patterns. Given the high orthogonality between isoforms discussed in Section 5, this suggests that certain isoforms from a single TF family might independently participate in similar regulatory circuits. To illustrate this, we focused on the subnetwork containing those isoforms (**Figure** 7C, **Figure** S7A), including the circuit between *EOMES*–*HMGA2*. *EOMES* had been broadly reported to be crucial for embryonic development of mesoderm^57–59^ and induction towards endoderm^60,61^, and *HMGA2* was reported to induce ectodermal differentiation with high expression^62^ and also involved in several early organogenesis process during gastrulation^62–64^. *EOMES*–*HMGA2* was also a typical case that suffered severe dropout in expression but showed clear negative correlation in TFA (**Figure** 7D, **Figure** S8B). In the network, we noticed that three *HMGA2* isoforms had negatively correlated relation with *EOMES-2*. Additionally, the pairs of *EOMES-2* versus *HMGA2-1*, *4* and *5* showed significant negative correlation among all three isoforms of *EOMES* and five of *HMGA2* (**Figure** 7E, **Figure** S8C).

Next, we tried to decipher characteristics that made these three *HMGA2* isoforms exhibit similar negative correlation with *EOMES-2*. Regulome of these isoforms only partially overlap, with their shared target genes also shared by the other two isoforms (**Figure** 7F) and exhibited low Jaccard indices among the five isoforms (**Figure** 7G), excluding the effect of same regulome leading to equal TFA. As we traced back to sequence level, i.e. transcript sequence, coding sequence (CDS) and amino acid sequence, the five *HMGA2* isoforms showed no sign of obvious cluster (**Figure** 7H, I, **Figure** S7D), suggesting that variation in function could be challenging to predict from sequence^28^. Analysis of *EOMES* holds the same viewpoint (**Figure** S7E).

To summarize, we applied our isoform-resolved regulome (CRED) and TFA pipeline to reconstruct bifurcation gene circuits in human gastrulation. The identified circuits were consistent with prior biological knowledge and uncovered new candidate regulators. More importantly, our analysis highlighted the functional convergence of distinct isoforms, even when their regulomes and sequences differ. These results underscore the need for TFA-based, isoform-level inference to decode gene regulatory logic in development.

**Figure 7.**
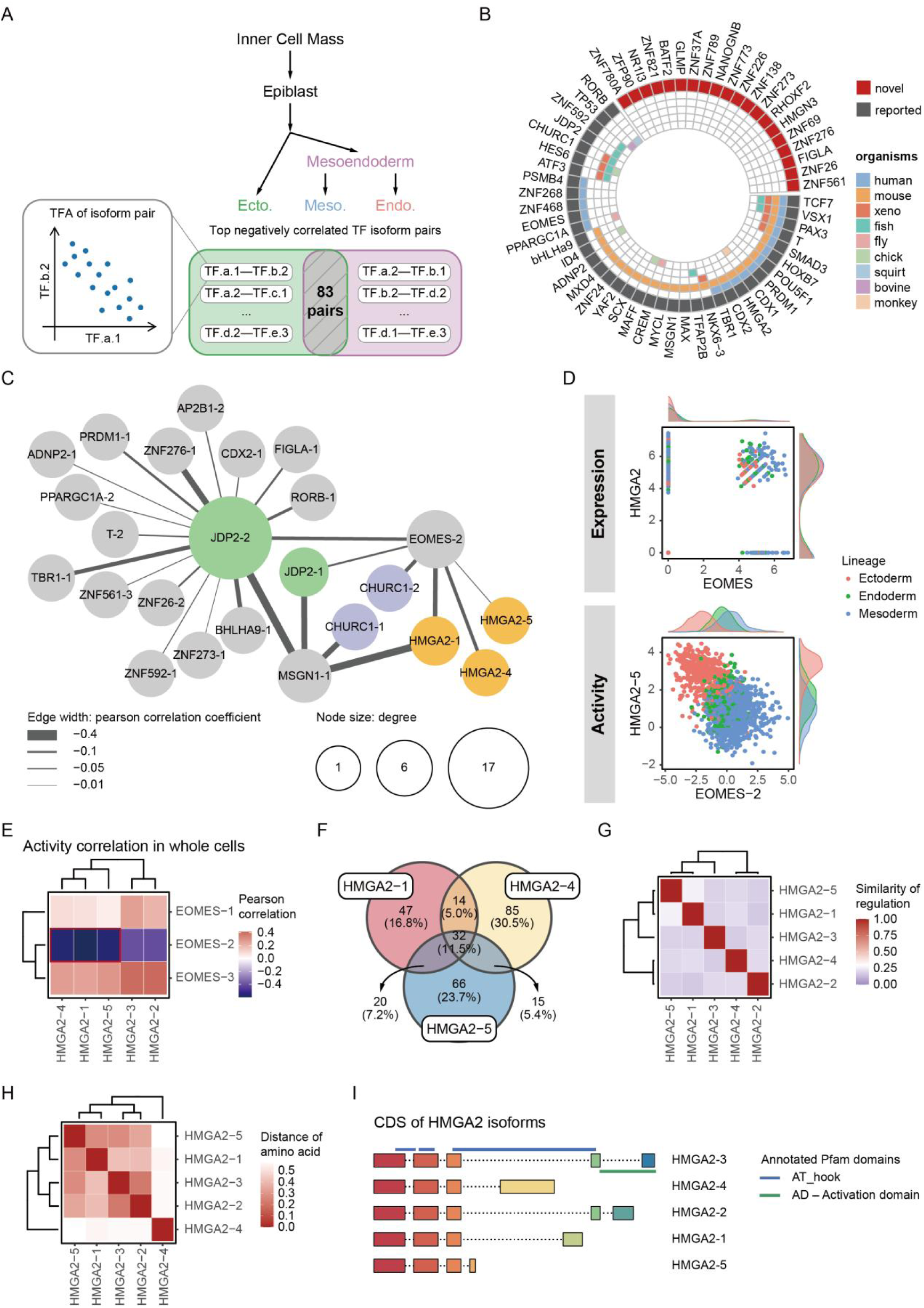
Isoform-level gene circuits in human gastrulation. (A) Schematic workflow for characterizing isoform-level gene circuits in human gastrulation. TF.x.n denotes No.n isoform of TF *X*. Region with gray shades contains 83 pairs, denoting overlap of isoform-level TF pairs top negative-correlative in their TFA in ectoderm cells and in mesoderm and endoderm cells together. Source data of all subplots in this figure is the same dataset in Figure 6. (B) TFs involved in the 83 pairs. Gray blocks annotate TFs that were reported to associate with gastrulation, where TFs in red characters show high TFA in germ layers that were not reported. Inner color blocks annotate organisms of literature reports. (C) Assembled isoform-level gene network using gene circuits in the 83 pairs containing different isoforms of one TF. Colored circles annotate isoforms of same TF. Node size denotes node degree. Edge width denotes the correlation of isoform pairs in mesoderm and endoderm cells, with wider edges reflecting stronger negative TFA correlations. The same network with edges denoting correlation in ectoderm is shown in **Figure** S7A. (D) Expression (top) and TFA (bottom) of *EOMES* and *HMAG2*. For isoform-level TFA, one representative isoform pair is shown here; additional pairs are provided in Figure S7C. (E) TFA correlation of all *EOMES* and *HMAG2* isoforms, calculated with whole cell population. Red square marks the pairs with significant negative correlation. (F) Regulatory targets overlap among three *HMAG2* isoforms. (G) Similarity of regulation of all *HMAG2* isoforms, calculated by Jaccard index. (H) Distance of amino acid sequence corresponding to all *HMAG2* isoforms, see **Methods**. (I) Coding sequence (CDS) diagram of all *HMAG2* isoforms with annotation of functional domains.

## Discussion

Transcription factors (TFs) sit at the top of gene regulatory networks. Despite often exhibiting low mRNA abundance, they exert massive influence by activating or repressing hundreds to thousands of downstream targets through direct binding and indirect regulatory cascades, thereby orchestrating entire cellular programs^65,66^. In this manner, interconnected circuits of TFs serve as critical drivers of cell fate decisions, ranging from lineage choices in development to state shifts in disease^67^. A pressing question in systems biology persists: Does inferred transcription factor activity (TFA) surpass direct mRNA abundance measurements in characterizing TF–TF regulatory circuits, and if so, where and how?

Our work says yes, especially in complex situations like developmental antagonism between TFs or tracing past regulatory “footprints” in end-point cell populations. Instead of simply estimating TFA across conditions, our work put forward the use of TFA to a more systematic view. Through *in silico* simulations (bipotent, unipotent and hematopoietic networks) and real-world datasets (spatial heart transcriptomics, and single-cell datasets of human hematopoiesis, gastruloids and preimplantation embryos), we showed that TFA consistently uncovers essential TF circuits in fate decisions, like *MESP1*–*MEOX2* exclusion where expression alone fails due to noise or dropouts, or traces the regulatory history of upstream mastering TF circuits, like *GATA1*–*SPI1* toggle which were mutual exclusively expressed in endpoint cell populations. At its core, this edge stems from TFA’s summation effect: it aggregates the collective behavior of downstream targets into a functional score, turning gene regulatory networks (GRNs) into integrated systems where net outputs emerge from direct and indirect interactions, much like weighted regulon enrichment in tools such as VIPER^21^ or DoRothEA^22^. In essence, TFA reinforces the identification of functioning TF circuits by capturing the propagated, top-down essence of GRN that expression misses, amplifying signals in networked biology—particularly through transited regulomes, which shine in noisy, dynamic developmental contexts because they embrace this non-linear summation rather than isolating linear bindings.

However, TFA’s advantage isn’t unconditional. Instead, it critically depends on regulome choices. Counterintuitively, transited (indirect) regulomes outperform direct ones in dynamic systems like development, which may stem from the summation effect we noted earlier: cell-fate-related circuits rely on cascades where master TFs exert upstream influence, rippling indirectly to endpoints via intermediaries, building hierarchical robustness^68^. Evolution favors such resilient setups, where summation buffers noise (e.g., dropouts) and amplifies signals, akin to gain in control theory^69^. Direct regulomes offer mechanistic purity and reflect the temporal function, but miss this propagation; transited ones, inferred from correlations or perturbations (like our Consensus REgulome Database (CRED) via overexpression), grasp system-level influence through network diffusion^70^, ideal for retrospective tracing—though they risk over-inference without benchmarking, as we did against DoRothEA^22^ and TRRUST^26^. Extending this, transited views could pinpoint therapeutic targets in TF-driven diseases like leukemia by capturing full ripple effects^71^.

A second key condition is that cell-type-specific regulomes outperform comprehensive, cross-tissue aggregates. Our humam embryo stem cell (hESC)-specific CRED database, derived from isoform overexpression in hESCs, exhibit better recall of active TFs in combinatorial perturbations and clearer germ-layer specificity in peri-gastruloids than regulome from merged resources. Biologically, we suggest this arises from TF context-dependency: binding sites and targets shift across cell types, shaped by chromatin accessibility, cofactors, and epigenetic states ^3,27^. Broad regulomes inject noise from mismatched contexts (e.g., cancer lines), weakening signals in stem cell niches. In ESCs, where pluripotency relies on tight regulatory wiring^72^, this specificity boosts TFA’s precision, illuminating why CRED excelled and why expansive databases don’t always equate to better ones. Yet, this condition also reveals a trade-off: while specificity sharpens fidelity, it impedes broader applicability. Here, we proposed a paradigm CRED in need of tissue-specific regulome whose extraction methodology can be adapted to broader contexts or species.

A third key condition is that isoform resolution proves essential, since gene-level views miss the functional divergence among variants. This arises biologically from alternative splicing, which yields isoforms differing in domains, localization, or binding affinity^27–30^, allowing precise tuning in development, such as switching at pivotal stages^55,56^. With CRED, we quantified this orthogonality, showing most isoforms govern unique target sets decoupled from broader TF roles. Subtly, this prevalence likely stems from evolutionary gains in regulatory diversity without gene duplication^28^. Overlooking it muddles signals, as in our *EOMES*–*HMGA2* circuit where isoforms converged on antagonism despite varied regulons and sequences. In turn, isoform-aware TFA exposes hidden modularity in GRNs, enhancing circuit reconstruction.

While these conditions empower TFA for robust isoform-level TF-circuits inference, limitations remain. Simulations, while insightful, rely on abstracted motifs that may overlook stochasticity or non-transcriptional layers like protein stability^13,14^. Transited regulomes risk over-abstraction, sacrificing mechanistic detail for breadth and potentially introducing artifacts from inferred edges. CRED’s reliance on overexpression data could amplify non-endogenous interactions, and TFA’s dropout handling assumes regulon resilience, yet severe noise might still bias results. Moreover, our ESC-centric focus limits generalizability to differentiated or adult tissues, where targeted regulomes still to be developed.

Looking ahead, experimental perturbations, such as isoform-level CRISPR knockouts in gastruloids, would validate circuits like *EOMES*–*HMGA2*. Extending beyond development, TFA could be employed to probe TF circuits in immune responses (e.g., inflammation toggles) or aging, where factors like NF-κB regulate senescence—raising questions on whether regulome specificity shifts with cellular aging, perhaps eroding orthogonality amid epigenetic drift^73,74^. Ultimately, by refining TFA under these conditions, our framework bridges computation and biology, advancing nuanced views of cell fate dynamics in health and disease.

## Supplementary information

Supplementary information includes Figure S1–7, Table S1 and Supplementary References, and can be found with this article online.

## Acknowledgements

We thank Y. Zhao for advice of algorithm applications on real biological processes; P. Zhou for insightful and generous feedback on data interpretation; J. Wu for providing original raw-data processing R scripts; L. Lambourne for visualization of alternative splicing; and the entire Zhiyuan laboratory for support and advice.

## Fundings

This work was supported by the National Key Research and Development Program of China (No. 2024YFA0919500, 2021YFA0910700) and National Natural Science Foundation of China (No. T2321001). Z.L. was supported in part by the Peking-Tsinghua Center for Life Sciences.

## Author contributions

Conceptualization, G.X.; Methodology, W.L., G.X. and Z.Z.; Software, W.L. and G.X., H.D. and Z.Z.; Formal Analysis, W.L., G.X. and H.D.; Investigation, W.L. and G.X.; Resources, W.L. and G.X.; Data Curation, W.L. and H.D.; Writing – Original Draft, W.L. and G.X.; Writing – Review & Editing, W.L., G.X., H.D., W.F., Y.Z., X.Y. and Z.L.; Visualization, W.L., G.X., and H.D.; Supervision, G.X. and Z.L.; Project Administration, G.X. and Z.L.; Funding Acquisition, Z.L.

## Declaration of interest

The authors declare no competing interests.

## Notes

### Competing Interest Statement

The authors have declared no competing interest.

