## Supplementary Information for "Graph-based tracing of dynamically functioning gene circuits in cell fate decisions with isoform resolution"

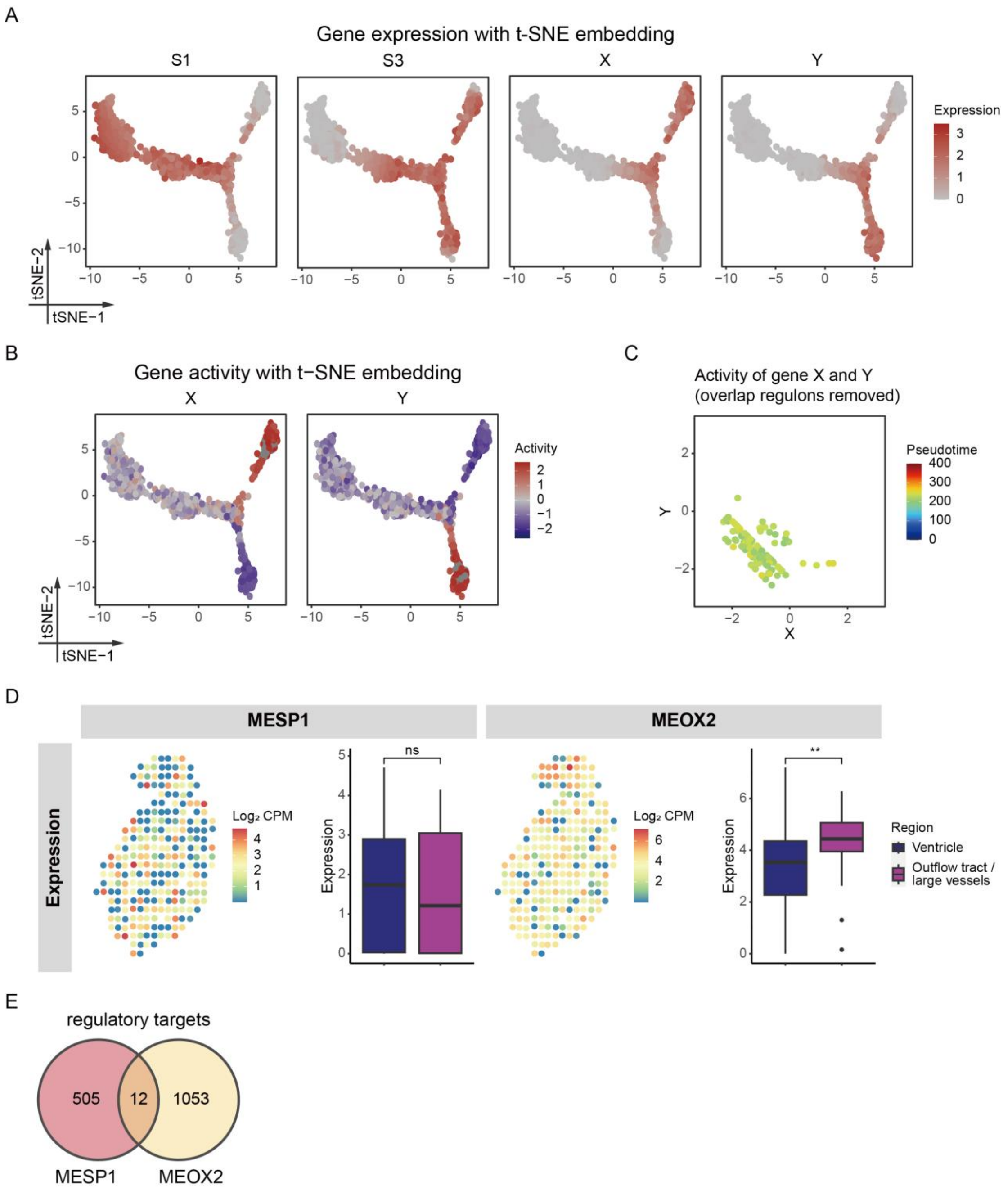


**Figure S1. Analysis of the Bipotent Network simulated data and spatial transcriptomic data from Asp et al.**[1]**.**

(A) Expression of *S1*, *S3*, *X* and *Y* in T-SNE plot. Source data of (A-C) is simulated result using the Bipotent Network in **Figure** 1B.

(B) TFA of *X* and *Y* in T-SNE plot.

(C) TFA of *X* and *Y* of intermediate samples in the range of 200-250 pseudotime, corresponding to lineage specification stage. Overlap regulons of *X* and *Y* were masked when calculating TFA.

(D) Expression of *MESP1* and *MEOX2* in spatial position and anatomical region. n.s.,

no significance, $p \geq 0.05$; $**$, $p < 0.01$. Source data of this subfigure is spatial transcriptomic data of human embryonic cardiac sample from Asp et al.[1].
(E) Overlap of *MESP1*'s and *MEOX2*'s regulatory targets. Source data of this subfigure is the merged regulome database of TRRUST[2] and DoRothEA[3].

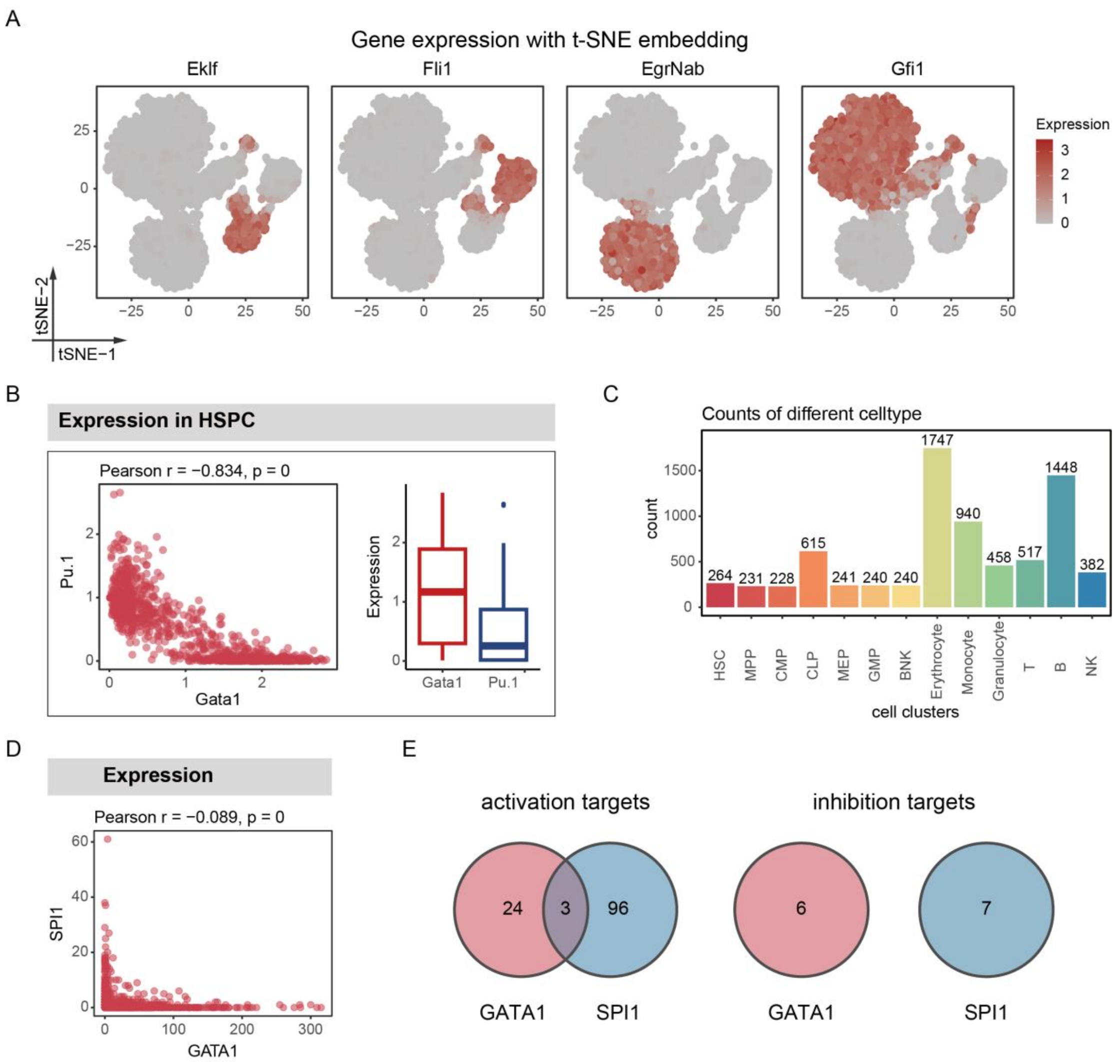


**Figure S2. Analysis of the 11 Nodes Hematopoiesis Network simulated data and Human blood cell atlas from Xie et al.[4].**
(A) Expression of *Eklf* (megakaryocyte marker), *Fli1* (erythrocyte marker), *EgrNab* (monocyte marker) and *Gfi1* (granulocyte marker) in T-SNE plot. Source data of (A) and (B) is simulated result using the 11 Nodes Hematopoiesis Network in **Figure** 2A.
(B) Expression of *GATA1* and *PU.1* in HPSC, hematopoietic progenitor / stem cell.
(C) Distribution of different cell types in Human blood cell atlas[4], source data of (D) is this dataset.
(D) Expression of *GATA1* and *SPI1* in Ery..
(E) Overlap of *GATA1*'s and *SPI1*'s regulatory targets. Source data of this subfigure is merged regulome database of TRRUST[2] and DoRothEA[3].

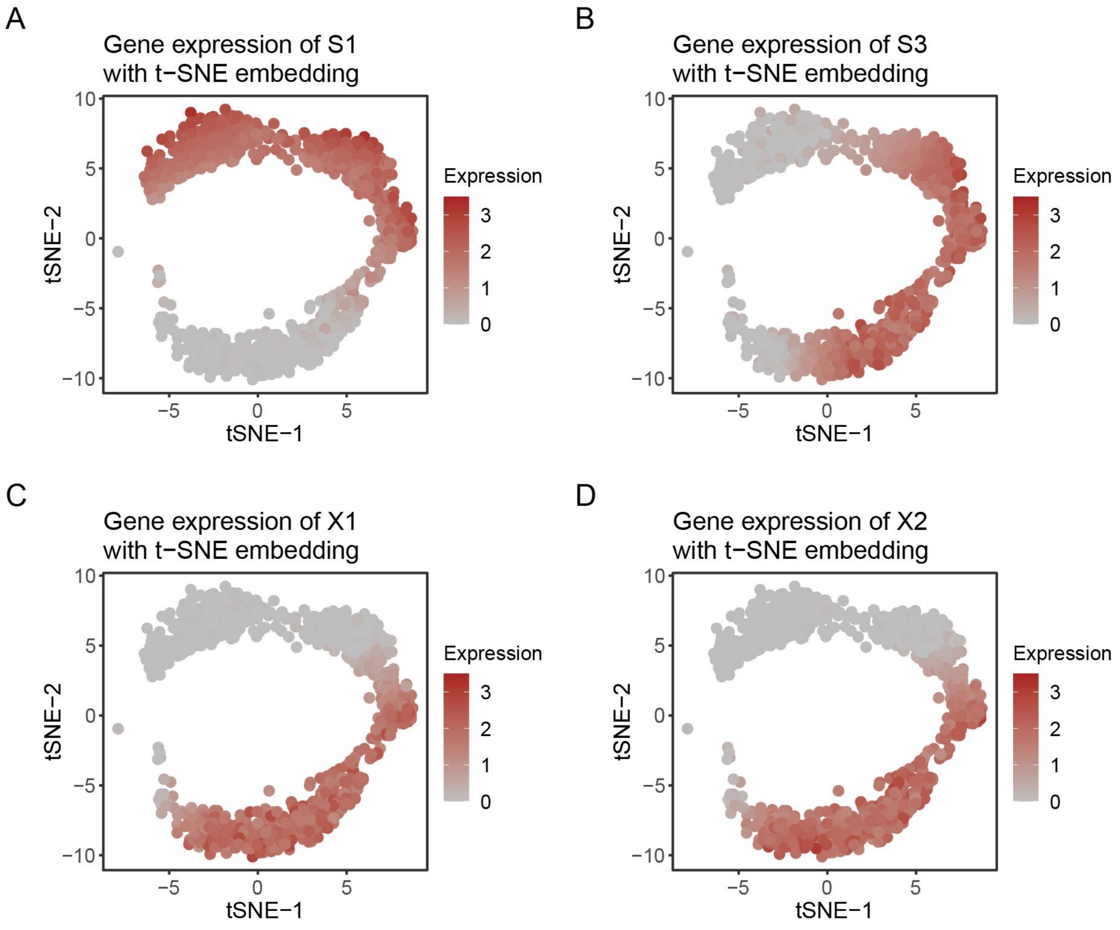


**Figure S3. Analysis of the Unipotent Network simulated data.**

(A) Expression of *S1* in T-SNE plot. Source data of (A-D) is simulated result using the Unipotent Network in **Figure** 3A.

(B) Expression of *S3* in T-SNE plot.

(C) Expression of *X1* in T-SNE plot.

(D) Expression of *X2* in T-SNE plot.

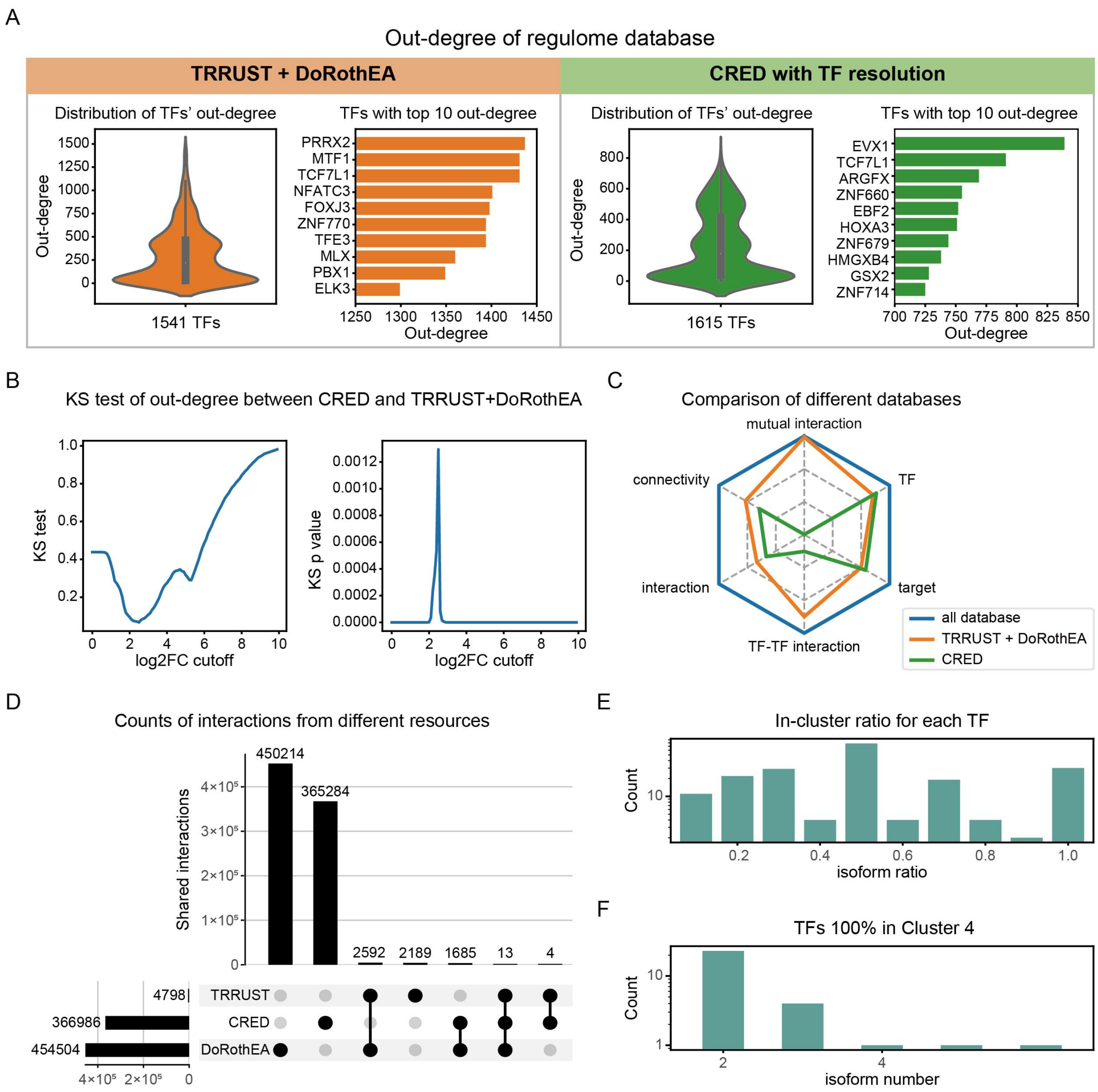


**Figure S4. Analysis of the CRED.**

(A) Distribution of out-degree and TFs with top out-degree in different regulome database.

(B) Kolmogorov-Smirnov test statistic of out-degree between CRED and merged database of TRRUST [2]and DoRothEA[3], calculated with gradient cutoff of log2 fold change (FC).

(C) Statistical comparison of different regulome databases.

(D) Counts and overlap of regulatory interactions from different resources.

(E) Distribution of the ratio of isoforms involved in Cluster 4 for each TF.

(F) Distribution of isform number for identical TFs (TFs with all isoforms in Cluster 4).

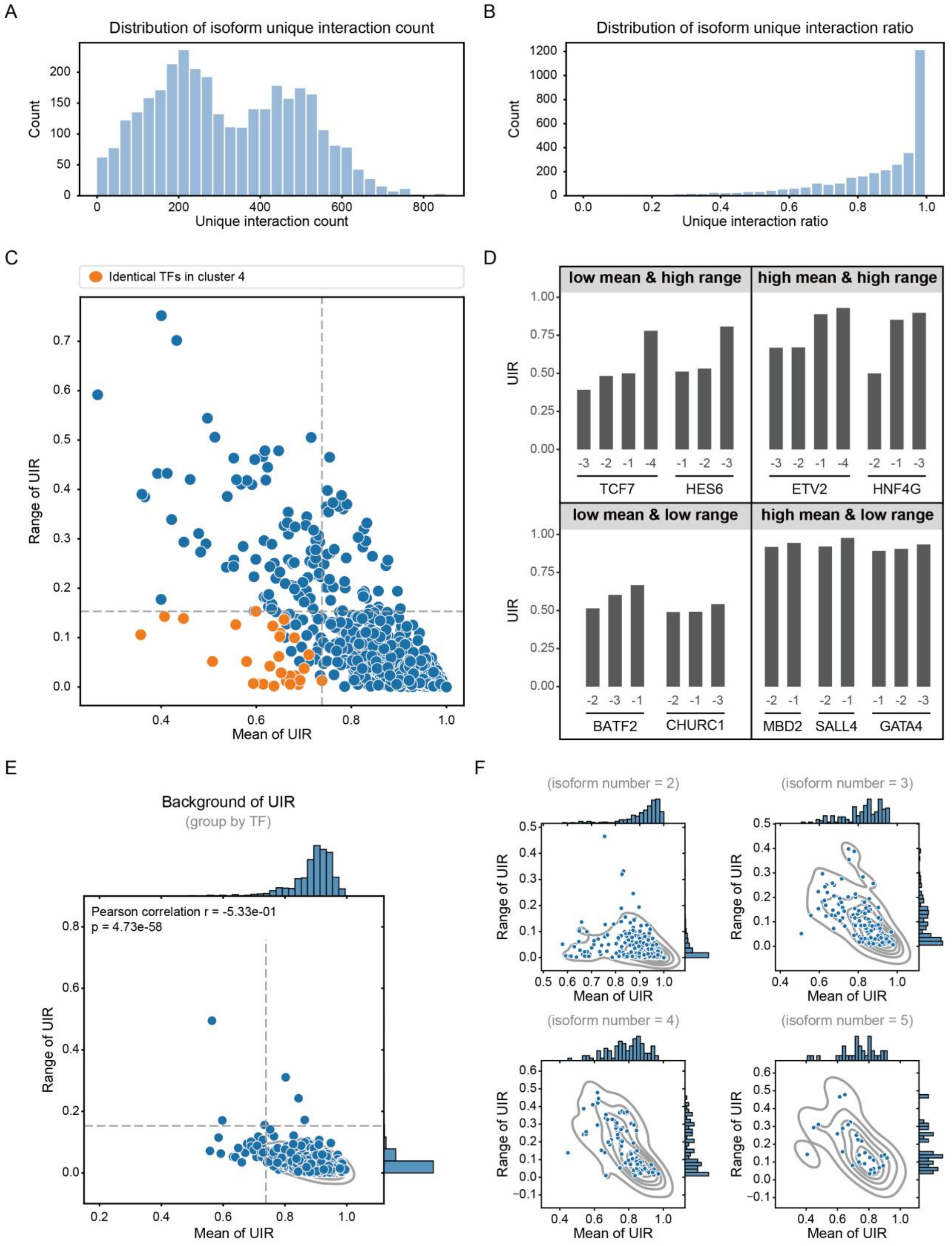


**Figure S5. Analysis of functional divergence among isoforms of the same TF**

(A) Distribution of isoform unique interaction counts.

(B) Distribution of isoform unique interaction ratio (UIR).

(C) Mean and range of UIR grouped by TF. Orange dots highlight identical TFs and dashed lines marked their maximum value.

(D) Typical cases of TFs with different level of mean and range of UIR.

(E) Mean and range of background UIR grouped for each TF with more than one isoform. Each dot represents one TF. Dashed lines represent thresholds defined by

"identical TFs" in (C). Calculation of background UIR see **Methods**.
(F) Mean and range of UIR grouped by TF, focusing on TFs with same number of isoforms.

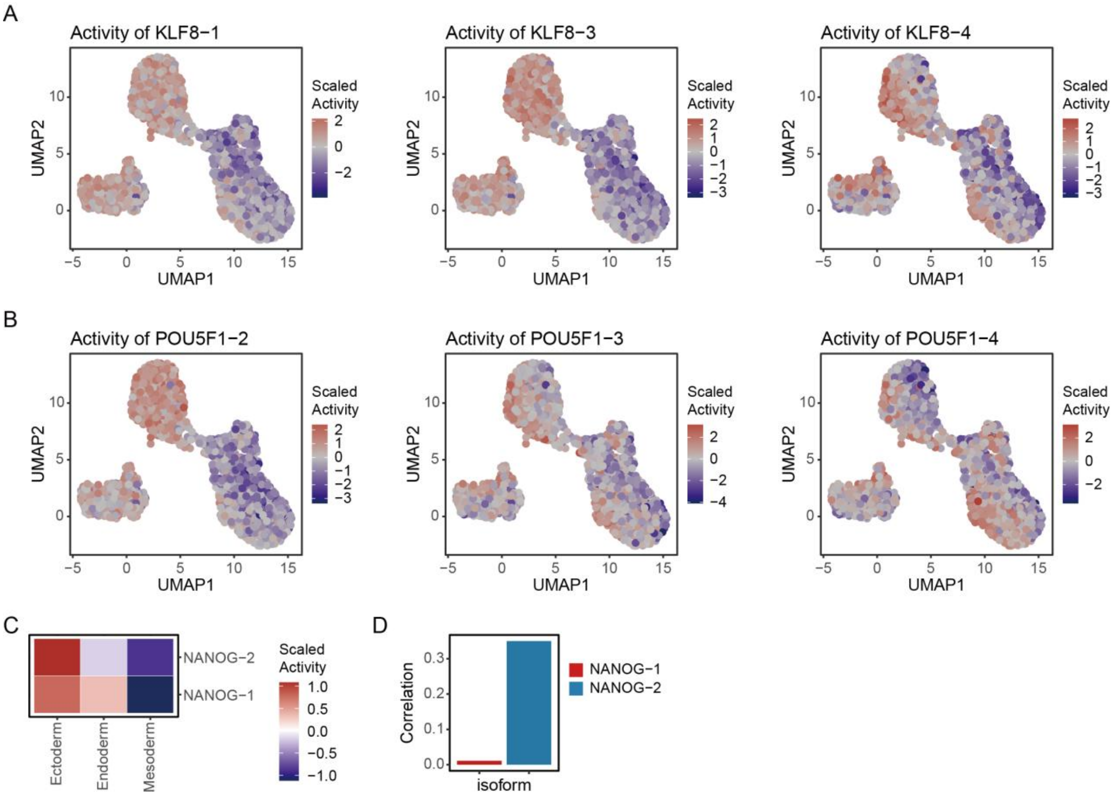


**Figure S6. Analysis of TFA in human peri-gastruloids from Liu et al.[5].**
(A) Scaled TFA of *KLF8* isoforms in UMAP plot. Source data of this figure is from Liu et al.[5].
(B) Scaled TFA of *POU5F1* isoforms in UMAP plot.
(C) Scaled TFA of *NANOG* isoforms cross three lineages.
(D) Correlation of expression and TFA of *NANOG* isoforms.

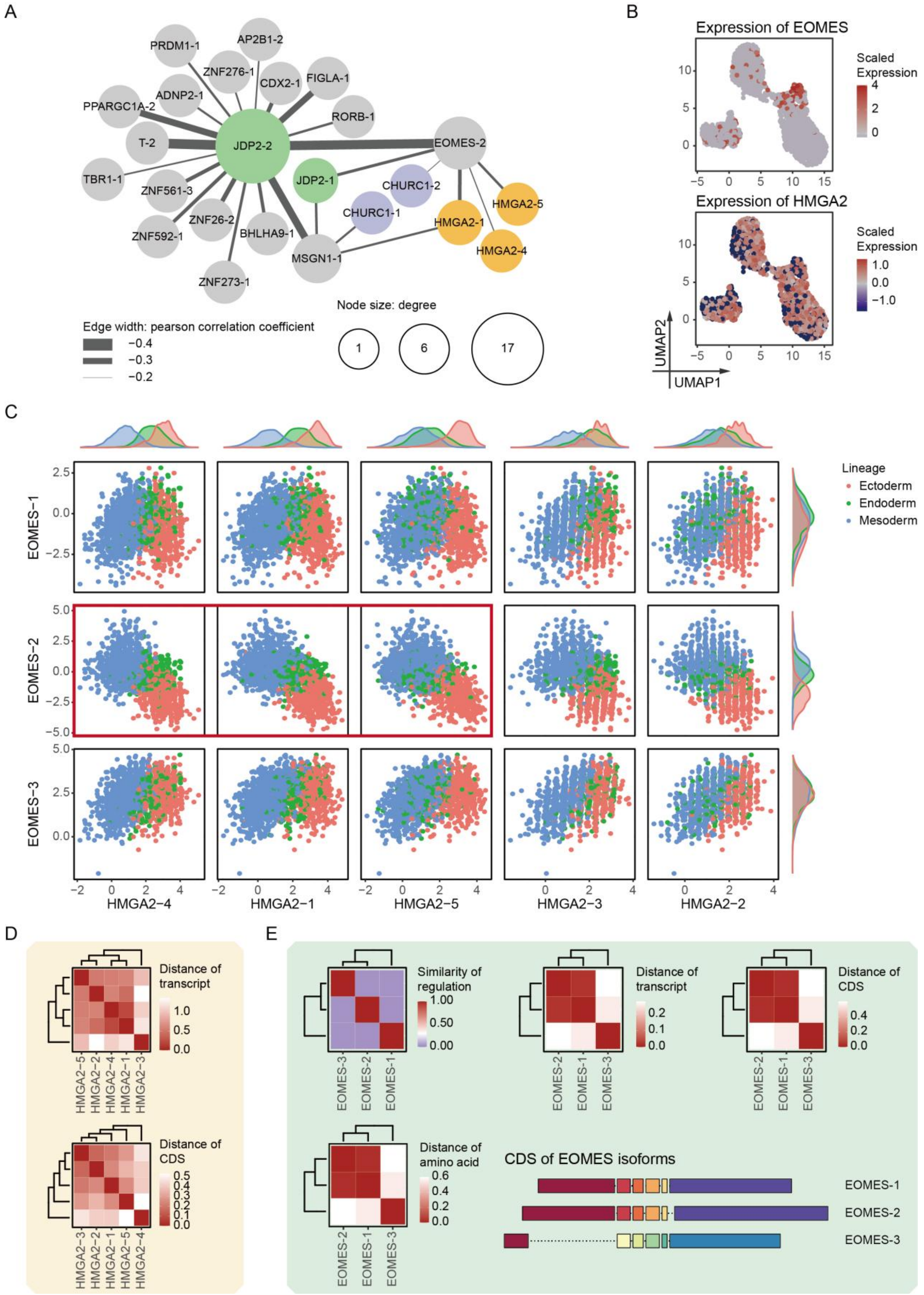


**Figure S7. Analysis of isoform-level gene circuits in human gastrulation.**

(A) Assembled isoform-level gene network using gene circuits in 83 pairs containing different isoforms of one TF. Colored circles annotate isoforms of same TF. Node size denotes node degree. Edge width denotes the correlation of isoform pairs in ectoderm cells, with wider edges reflecting stronger negative TFA correlations. The same network with edges denoting correlation in mesoderm and endoderm is shown in

**Figure** 7C.
(B) Scaled expression of *EOMES* and *HMGA2* in UMAP plot.
(C) Paired activity of isoforms of *EOMES* and *HMAG2*. Red square corresponds to that in **Figure** 7E.
(D) Distance of transcript sequence and coding sequence (CDS) corresponding to all *HMAG2* isoforms.
(E) Clustering analysis of all isoforms of *EOMES*, across regulon and multiple dimensions of sequence (transcript, CDS and amino acid).

**Table S1. Statistics of different regulome databases.**

| Database | Regulator | Target | Protein | Interaction |
|---|---|---|---|---|
| CRED (isoform-resolved) | 3250 | 23957 | 24350 | 1180085 |
| CRED (TF-resolved) | 1615 | 21954 | 22474 | 366986 |
| TD | 1541 | 20314 | 20320 | 456697 |
| TDC | 1911 | 30465 | 30478 | 821981 |

| Database | Activation | Inhibition | Connectivity | TF–TF interaction |
|---|---|---|---|---|
| CRED (isoform-resolved) | 1130532 | 49553 | 363.103077 | 45072 |
| CRED (TF-resolved) | 348962 | 18024 | 227.095297 | 14178 |
| TD | 411803 | 44894 | 296.364049 | 69342 |
| TDC | 759219 | 62762 | 430.131345 | 83291 |

| Database | TF–TF Mutual interaction | TF–TF Mutual activation | TF–TF Mutual inhibition |
|---|---|---|---|
| CRED (isoform-resolved) | 28021 | 25168 | 2853 |
| CRED (TF-resolved) | 48 | 34 | 4 |
| TD | 25262 | 21030 | 198 |
| TDC | 25516 | 21262 | 204 |

CRED: regulome database integrated in this work.
TD: established database represented by TRRUST[2] and DoRothEA[3].
TDC: integration of TF-resolved CRED and established database represented by TRRUST[2] and DoRothEA[3].
