## Supplementary Method for "Graph-based tracing of dynamically functioning gene circuits in cell fate decisions with isoform resolution"

### Methods

**Calculation of TFA**

For TFA calculation, we employed a published algorithm VIPER (virtual inference of protein activity by enriched regulon analysis)[1] based on its consistency with our theoretical framework (**Figure** 1A). VIPER employs a probabilistic framework to infer TFA from gene expression profiles by computing rank-based normalized enrichment score, utilizing the prior knowledge of regulome. To meet the need of single-cell analysis in our work, we used single sample version function 'viper' in R package viper and chose 'ttest' to compute gene expression signatures. The parameter 'minsize' (the minimum number of targets allowed per TF) was set to 4 by default, while it would be lowered for small GRNs.

To estimate differential TFA of single sample treatment group against multiple sample control group, we developed a optimal version called 'viper_enhanced'. In computation of gene expression signature, we refined the algorithm by comparing the treatment sample against control samples using the Student's t-test, rather than comparing it with all other samples.

For choice of regulome, we used:

1. **Manually designed regulons**, derived from given GRN, for analysis of the Bipotent Network, the 11-Node Hematopoiesis Network and the Unipotent Network;
2. **TD**, merged database using TRRUST[2] and DoRothEA[3] representing current knowledge of regulatory network, for analysis of spatial transcriptomic data of developing human heart[4], atlas of human blood cells[5], and comparison analysis among regulome databases of combinatorial TF overexpression in human embryonic stem cells (hESCs)[6] and human stem-cell-derived peri-gastruloids[7];
3. **CRED**, regulome database with isoform resolution integrated in this work, for analysis of human stem-cell-derived peri-gastruloids[7], using consensus regulome with TF resolution for comparison analysis among regulome databases of combinatorial TF overexpression in hESCs[6] and human stem-cell-derived peri-gastruloids[7];
4. **TDC**, merged database using TRRUST[2], DoRothEA[3] and TF resolution CRED, for comparison analysis among regulome databases of combinatorial TF overexpression in hESCs[6].

In this work, regulatory interactions we aimed to testify would be masked in corresponding analysis. Any dataset-specific settings would be described within corresponding dataset description.

**Analysis of single-cell level simulation based on GRN**

To investigate feasibility of TFA in simplified systems, we performed several simulations and analyzed through the following steps:

**Step 1: GRN construction**

In this work, we constructed three representative GRNs: the Bipotent Network (**Figure** 1B, a basic GRN that regulates cell fate decisions between two antagonistic fates), the 11-Node Hematopoiesis Network (**Figure** 2A, a well-documented GRN for myeloid differentiation in hematopoiesis[8]) and the Unipotent Network (**Figure** 3A, a GRN that regulates cell state transition from unipotent progenitors to mature cells and gradually activation of downstream gene programs, in this work we set 5 genes in each module). For regulatory logics among regulations towards the same target, the 11-Node Hematopoiesis Network followed the boolean regulation rules displayed by Krumsiek et al.[8]. The other two networks followed the principle of activation in OR logic and inhibition in AND logic, that is, the target gene would express while at least one of its activation condition and none of its inhibition condition exists.

**Step 2: Single-cell level simulation**

We used Python package BoolODE[9] to generate single-cell transcriptome data with stochastic non-linear ordinary differential equation (ODE) based on input boolean GRN and initial cell state.

For the Bipotent Network, the initial cell state was set with 1 unit exclusive expression of gene *S1*. Simulation was performed 2,000 rounds with a maximum time of 800, sampling one cell from each simulation, while we further restricted the time window to 0–400 representing the procedure of priming state, cell fate specification and cell fate commitment, remaining 1,010 cells. 125 cells at time window 200–250 were defined as intermediate samples.

For the 11-Node Hematopoiesis Network, the initial cell state was set with 1 unit expression of *Cebpa*, *Pu.1* and *Gata2* as recommended. Simulation was performed 8,000 rounds with a maximum time of 800, sampling one cell from each simulation.

For the Unipotent Network, the initial cell state was set with 1 unit exclusive expression of gene *S1*. Simulation was performed 2,000 rounds with maximum time of 800, sampling one cell from each simulation, while we further restricted the time window to 0–500 representing the transition from progenitor state to mature state, remaining 1,265 cells.

**Step 3: TFA calculation**

TFA calculation was performed as detailed in previous section.

For the Bipotent Network, we masked the known mutual inhibition of gene *X* and *Y*, as we aimed to infer regulatory relationship between the two genes. Further, we masked overlapping regulons to decouple the inherent relationship between *X* and *Y*, i.e., removing *X*'s inhibition to *Y1*–*Y5* and *Y*'s inhibition to *X1*–*X5*.

For the 11-Node Hematopoiesis Network, we masked the known mutual inhibition of *Gata1* and *Pu.1*, as we aimed to capture this functioning circuit from the endpoint cell population.

For the Unipotent Network, we defined transited regulons, i.e., *S3*'s indirect activation to module1 and module2. TFA was calculated with two sets of regulome, one containing only direct regulations and the other including transited regulons.

**Step 4: analysis and visualization**

T-Stochastic Neighbor Embedding (T-SNE)[10] dimension reduction was performed by R package Rtsne and Seurat[11]. For the 11-Node Hematopoiesis Network, cell clusters were defined by function 'FindClusters' in R package Seurat[11] and then annotated by gene expression. Results were visualized by R package ggplot2[12].

**Analysis of public datasets**

To testify usage of TFA in real-world systems, we analyzed several public datasets in this work.

**Spatial transcriptomic data of developing human heart[4]**

Raw count matrix of a dataset of spatial transcriptomic data of developing human heart was downloaded from https://www.spatialresearch.org. The dataset was generated using the Spatial Transcriptomics v1.0 protocol and annotated as four anatomical regions (ventricle, cluster 0-3; atrium, cluster 4; outflow tract/large vessel, cluster 5; and other, cluster6-9). We preformed gene expression imputation for each spot using Tangram[13] with corresponding single-cell RNA-seq data as reference, following the methods in a previous work[14]. Raw counts were normalized to counts per million (CPM) and then log-transformed ($log_2$(CPM+1)). We exemplified a tissue section with most detected spots (section 16, 243 spots) from this dataset, collected at 9 post-conception weeks (**Figure** 1F). Using R package viper with regulome TD, activity profile including 1,370 TFs was calculated. To nominate region-exclusive TF pairs in this dataset, we did t-test between activities in the two main anatomical regions ventricle and outflow tract/large vessels for all TFs and ranked them by t-test statistics. TFs with top ranks in the two regions respectively would be selected (In this work, we selected *MESP1*, ranking 3/1,370 in ventricle, and *MEOX2*, ranking 4/1,370 in outflow tract/large vessels, as showcasing example). Results were visualized by R package ggplot2[12], ggpubr and gridExtra.

**Atlas of human blood cells**[5]

Single-cell transcriptome data and metadata including cell type annotations of 7,551 human blood cells were downloaded from web portal (http://scrna.sklehabc.com/). Raw counts were normalized to counts per million (CPM) and then log-transformed ($log_2$(CPM+1)). For TFA calculation, we utilized regulome TD and masked mutual inhibition of *GATA1* and *SPI1* that was widely documented to regulate myeloid differentiation. Using R package viper with regulome TD, activity profile including 1,388 TFs was calculated. Results were visualized by R package ggplot2[12], ggpubr and gridExtra.

**Combinatorial TF overexpression in hESCs**[6]

SHARE-RNA-seq of combinatorial TF overexpression in hESCs was obtained from the GEO with accession code GSE217066. In this dataset, hESCs were transducted with 10 TF open reading frames (ORFs) in combinations and cultured for 7 days. The subsampled dataset containing 30,468 single-cells was grouped by combinations (including 9 single TFs (TF+GFP control), 35 double TFs and 2 triple TFs) and generated into pseudo bulk profiles by mean expression. Control samples (n=1000) expressing GFP from another dataset in the same work were obtained from the GEO with accession code GSE217460, and generated into 3 pseudo bulk profiles by mean expression, each containing 50% cells. Raw counts were normalized to counts per million (CPM) and then log-transformed ($log_2$(CPM+1)). Genes with top 80% expression in each combination sample and detected in control group (expression>0 in at least one pseudo bulk control profile) were used to calculate TFA respectively. Here we used function 'viper_enhanced' optimized in this work to calculate differential TFA against control group to magnify effect of TF overexpression. Recall rate was defined as the proportion of TFs in combinations whose TFA exceeded percentage rank threshold, calculated with regulome TD, CRED with TF resolution, and TDC. Results were visualized by R package ggplot2[12].

**Human stem-cell-derived peri-gastruloids**[7]

Single-cell RNA-seq data of day 11 peri-gastruloids was obtained from the Gene Expression Omnibus (GEO), with accession code GSE232861. From the total of 10,684 cells, we selected those belonging to the three germ layers (annotated as Epi, Ect, PS, Nas.meso, Adv.meso, DE, Extra.endo and ExE.meso), then randomly downsampled to 20% of this cell population, remaining 1,930 cells for the following analysis in this work. Python package Scanpy[15] was used to perform normalization by reads per cell and log normalization on the raw data, identify 2,000 highly variable gene and visualize using Uniform manifold approximation and projection (UMAP)[16]. Using R package viper (parameter 'minsize' = 2), we calculated TFA on this dataset with three regulome databases (For regulome TD, activity profile including 1,457 TFs was calculated. Regulome CRED with isoform resolution, 3,237 isoforms. Regulome CRED with TF resolution, 1,566 TFs.).

In the screening of the Human TFome, a group of 65 TFs was identified to induce differentiation individually, in context of three human induced pluripotent stem cell (iPSC) lines[17]. We exemplified this group of TFs as high activity TFs detected in context of the three germ layers and calculated recall rate in the dataset of human stem-cell-derived peri-gastruloids. Recall rate was defined as the proportion of differentiation-inducing TFs whose TFA exceeded percentage rank threshold, calculated with regulome TD and CRED with TF resolution, along with corresponding background model (randomly shifting ranks of TFs).

To extract isoform-level gene circuits in human gastrulation, we calculated TFA correlation for all isoform-level TF pairs in ectoderm cells and mesoendoderm cells. Top 2500 negatively correlated pairs in ectoderm and mesoendoderm were selected respectively, remaining 83 overlapping pairs. The network containing those isoforms was visualized with Cytoscape.

For literature searching of defining novel regulators, the symbol name of each unique TF involved in 83 pairs (n = 58) along with each germ layer were queried in 4 different searching engines on March 30, 2024. Searching was performed in Bing for the first round, then in PubMed and in Google Scholar, and finally in GPT-4, making sure no related paper was leaked. TFs were considered ‘reported’ if at least one paper stated that the TF is preferentially expressed or functions as marker in the germ layer consistent with where it showed high TFA in our analysis; otherwise, the TF was considered ‘novel’. Literature based on classical model organisms were considered. The novel and reported TFs and the supporting species were illustrated through a circular heatmap using R package circlize[18] and ComplexHeatmap[19].

Visualization not mentioned above was performed by R package ggplot2[12], ggExtra, ggpubr, ggsci and pheatmap.

**Human preimplantation embryo**[20]

Isoform-resolved transcriptome of the human preimplantation embryo was obtained from the GEO with accession code GSE190547. This dataset contained 73 human embryos representing six embryonic developmental stages: zygote (1C, n=13), 2-cell (2C, n=13), 4-cell (4C, n=16), 8-cell (8C, n=15), morula (n=3) and blastocyst (n=13). Raw counts were normalized to counts per million (CPM). Isoforms were aligned by Ensembl Transcript ID. Results were visualized by R package ggplot2[12].

**Integration of current widely used regulome databases**

We integrated regulome TD containing 456,698 TF-target interactions for 1,541 human TFs and their mode of regulation (MoR, activation or repression) from current widely used regulome databases, TRRUST[2] and DoRothEA[3] following methods from our previous work[21]. DoRothEA is a built-in collection of human regulome in R package DoRothEA (version 1.8.0) including 454,504 TF-target interactions for 1,541 human TFs. TRRUST is a manually curated database of transcriptional regulatory networks, whose latest version contains 8,427 TF-target regulatory relationships of 795 human TFs with annotation of MoR. Here, interactions with unknown MoR were excluded. For interactions showing conflict MoR in the two resources, TRRUST was taken as more credible. However 207 interactions had conflict MoR records in TRRUST, then the interactions would be retain with its MoR as recorded in DoRothEA or be omitted if absent in

DoRothEA.

**Extracting Curated REgulome Database (CRED)**

For extraction of regulome with isoform resolution, we utilized a TF Atlas of directed differentiation that overexpressed 3,548 isoforms of 1,836 TFs respectively in hESCs[6]. The filtered and downsampled single-cell RNA-seq data was obtained from the GEO with accession code GSE217460, containing 671,453 cells overexpressing 3,266 isoforms of 1,714 TFs. Differentially expressed genes (DEGs) of cells overexpressing each isoform and control cells expressing GFP were calculated with funciton 'sc.tl.rank_genes_groups' in Python package Scanpy[15] (method=wilcoxon, tie_correct=True). Here, genes with zero expression and the overexpressing TF itself were omitted.

For selection of expression fold change (FC) cutoff of DEGs, we made use of established database (represented by regulome TD). Due to the restriction that established regulome database only have TF resolution, we remained consensus regulations among isoforms of the same TF. To be specific, only genes that were up-regulated ($\log_2 FC>0$) in overexpressing all isoforms of the TF were considered as its activation targets, and *vice versa*. In screening of optimal cutoff of $\log_2 FC$, we restricted adjusted P-value to smaller than 0.05. For stepwise-screened cutoff, consensus regulations with mean $\log_2 FC$ greater than the cutoff were retained. Then, we performed Kolmogorov-Smirnov (KS) test with function 'stats.ks_2samp' in Python package scipy, between number of regulations remained for each TF and out-degree of each TF in regulome TD. Cutoff with minimum KS-test statistic was selected ($\log_2 FC$ cutoff=2.5). The remained consensus interactions were integrated as CRED with TF resolution. DEGs of overexpressing each TF isoforms that reach the same threshold ($\log_2 FC$ cutoff=2.5, adjusted P-value<0.05) were integrated as CRED with isoform resolution.

For integration of regulome TDC, we merged CRED with regulome TD and considered CRED as the most credible. That is, all interaction presented in CRED would be preserved, with CRED serving as a reliable reference for resolving any conflicting relationships.

Visualization was performed by Python package matplotlib.

**Analysis of regulatory similarity between isoforms**

For all isoforms in CRED (n=3250), regulatory similarity was quantitatively accessed with Jaccard index of their regulatory targets pair-wisely:

$$\text{Jaccard\_index}_{\text{isoform i, j}} = \frac{\text{number of identical regulons between i and j}}{\text{total of regulatory targets for i and j}}.$$

Isoforms were clustered and visualized by Jaccard index using function 'sns.clustermap' in Python package seaborn (threshold=1.6).

For detailed analysis of cluster 4, in-cluster ratio was defined as the proportion of isoforms in cluster 4. TFs with multiple isoforms and 100% in-cluster ratio were defined as identical TFs. We used R package clusterProfiler and database 'org.Hs.eg.db' to perform gene ontology (GO) enrichment analysis of biological process terms. Significantly enriched GO terms for identical TFs were filtered using Benjamini-Hochberg (BH) correction with an adjusted P-value<0.05 and a minimum gene count threshold of 3. We randomly generated 50 lists of TFs with the same size (n=30) and performed enrichment analysis on the same set of terms as background. Terms with gene ratio in identical TFs higher than upper quartile of background distribution were retained.

Only one of the terms consist of the same set of TFs was presented in visualization. Results were visualized by R package ggplot2[12].

**Analysis of UIR of isoform regulome**

Regulons that were unique in target or mode of regulation among regulome of all isoforms of each TF was defined as unique regulons (Fig. 5A). Unique interaction ratio (UIR) of isoform $i$ was defined as follows:

$$UIR_i = \frac{k_i}{N_i},$$

where $k_i$ denoted the number of unique regulons of isoform $i$ and $N_i$ denoted the total number of regulome of isoform $i$.

To estimate the background of UIR, we generated a random regulome by shuffling isoform-target pairs in CRED 5 times. Then, background UIR was calculated as its definition.

Results were visualized by Python package matplotlib, seaborn and R package ggplot2[12], ggpubr.

**Comparison among isoforms of same TF**

In this work, we exemplified HMGA2 and EOMES for clustering analysis across TFA, regulon and multiple dimension of sequence (transcript, CDS and amino acid). Clustering of TFA was performed on a downsampled set of 1,930 cells in the dataset of human stem-cell-derived peri-gastruloids[7]. For regulon overlap analysis, we utilized CRED with isoform resolution. For sequence analysis in each level (transcript, CDS and amino acid), alignment was performed with Matlab function 'multialign' and distance was calculated with Matlab function 'seqpdist', both with default parameters. Results were clustered and visualized with R package ggplot2[12], ggtree, aplot and ggvenn. The CDS diagrams with domain annotations were plotted following Python code from Lambourne et al[22].
